# Natural Language Prompts Guide the Design of Novel Functional Protein Sequences

**DOI:** 10.1101/2024.11.11.622734

**Authors:** Nikša Praljak, Hugh Yeh, Miranda Moore, Michael Socolich, Rama Ranganathan, Andrew L. Ferguson

## Abstract

The advent of natural language interaction with machines has ushered in new innovations in text-guided generation of images, audio, video, and more. In this arena, we introduce **Bio**logical **M**ulti-**M**odal **M**odel (**BioM3**), as a novel framework for designing functional proteins via natural language prompts. This framework integrates natural language with protein design through a three-stage process: aligning protein and text representations in a joint embedding space learned using contrastive learning, refinement of the text embeddings, and conditional generation of protein sequences via a discrete autoregressive diffusion model. BioM3 synthesizes protein sequences with detailed descriptions of the protein structure, lineage, and function from text annotations to enable the conditional generation of novel sequences with desired attributes through natural language prompts. We present *in silico* validation of the model predictions for subcellular localization prediction, reaction classification, remote homology detection, scaffold in-painting, and structural plausibility, and *in vivo* and *in vitro* experimental tests of natural language prompt-designed synthetic analogs of Src-homology 3 (SH3) domain proteins that mediate signaling in the Sho1 osmotic stress response pathway in baker’s yeast. BioM3 possesses state-of-the-art performance in zero-shot prediction and homology detection tasks, and generates proteins with native-like tertiary folds and wild-type levels of experimentally assayed function.

## 1 Introduction

Millions of years of evolution have shaped the diversity of extant protein sequences through mutation and selection to encode precise three-dimensional structures and biological functions and map primary sequences to multifaceted phenotypes, including foldability, biochemical activities, specificity, and organismal fitness in natural biological contexts [1, 2, 3, 4, 5]. These functional characteristics are encoded as patterns within the amino acid sequences. With the explosion of sequence data and computational resources, deep generative models have emerged as a powerful tool to learn these design principles and use these rules to generate novel proteins with controlled phenotypic properties. As these models learn the underlying design rules, our capacity to generate new hypotheses for protein mechanisms and design variants with novel sequences and functions dramatically improves, opening avenues in medicine, biotechnology, chemical engineering, and public health [6, 7, 8, 9]. Indeed, by learning from evolutionary patterns, both structure-based and sequence-based protein generative models have been shown to generate novel sequences with enhanced protein function [10, 11, 12, 13], produce libraries enriched with functional variants [14, 15, 16, 17], and enable design of sequences with engineered structure and function [18, 19, 20, 21]. A relatively untapped resource in this endeavor is the extensive corpus of textual annotations accompanying protein sequences within large databases that provide natural language descriptions of lineage, properties, and function. This raises the intriguing possibility that the rich information within this literature that could further inform deep generative models and complement protein-based models to enhance our understanding of protein function and guide protein design through natural language prompting.

Integrating natural language processing (NLP) with protein language models (pLMs) presents a means to harness both evolutionary sequence information and human knowledge embedded in protein sequence annotations. ProtST [22] was the first model to align protein and text representations, demonstrate the feasibility of joint embedding spaces, and accomplish downstream prediction tasks, while ProteinDT [23] was the first to leverage these joint representations to guide novel protein sequence generation and conduct *in silico* validation of the model predictions. Our framework, Biological Multi-Modal Model (BioM3), builds upon these pioneering studies by scaling the training corpus to 45M text-protein pairs, enriching text prompts with detailed family descriptions, and introducing homologous relationships within the joint embedding space. The model also incorporates a novel conditional autoregressive diffusion model that permits order-agnostic in-painting, expanding on EvoDiff [24] by introducing text-prompt conditioning of sequence generation. This informs a three-stage process comprising alignment of biomedical language models with protein language models, refinement of these embeddings, and text-guided generation of protein sequences (Fig. 1A). Since BioM3 has been trained over a vast corpus of protein sequences drawn from thousands of different families, it represents a generic and transferable model capable of text-guided generative design of protein sequences with diverse structure and function. Moreover, through the alignment of protein sequence and text annotations within a joint latent space, the model can also locate the closest annotations within the joint embedding for a particular protein sequence in order to perform protein sequence annotation and property prediction. We present *in silico* validations of the model in applications to subcellular localization prediction, reaction classification, remote homology detection, scaffold in-painting, and structural plausibility. We also conduct *in vivo* and *in vitro* assays of designed analogs of Src-homology 3 (SH3) domain proteins that mediate signaling in the Sho1 osmotic stress response pathway in baker’s yeast (Fig. 1B). To the best of our knowledge, this represents the first experimental validation of functional proteins designed using natural language text prompts. The integration of NLP and pLMs opens new frontiers in protein engineering, enabling intuitive and flexible protein design through text prompts, democratizing the process of functional process design, and paving the way for advances in synthetic biology and biotechnology.

**Figure 1:**
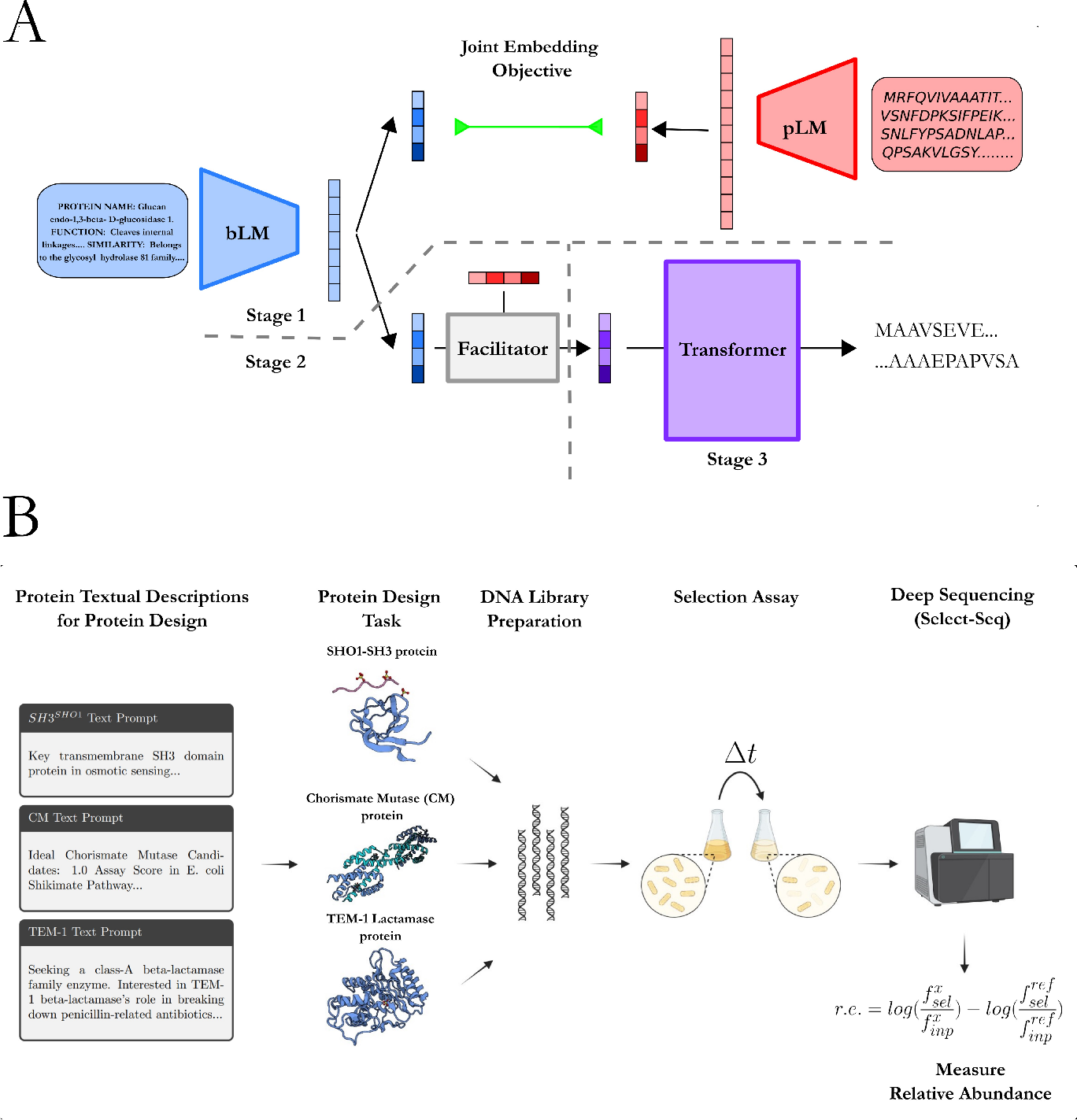
Overview of the BioM3 framework for protein design via natural language prompts. (A) The framework consists of three stages: (1) integration of a biomedical Language Model (bLM) and a protein Language Model (pLM) into a joint latent embedding space using Protein embeddings with natural language using Contrastive Learning (PenCL), (2) refinement and enhancement of the text embeddings (Facilitator), and (3) generation of novel protein sequences via text-conditioned autoregressive discrete diffusion (ProteoScribe). Training is performed over a corpus of 45M text-protein pairs. (B) The trained model is used to design protein sequences based on natural language text prompts providing textual descriptions for the lineage, properties, and/or function of desired target proteins. The generated protein sequences are subjected to *in vivo* and *in vitro* experimental validation via gene synthesis, high-throughput selection assays, next-generation sequencing, and biochemical binding assays to measure fitness and phenotype. Portions of this image were created in BioRender. BioRender.com/r83g055.

## 2 Related Work

### Protein Representation Learning

Protein Language Models (pLM) have played a pivotal role in learning effective protein representations for downstream applications in predictions of structure and function and understanding evolutionary lineage and design rules [25, 26, 27, 16, 28, 11]. The predominant training objectives for these pLMs are self-supervised learning tasks, notably masked token prediction [25, 29, 30], a technique borrowed from successes in natural language processing with BERT-based architectures [31]. Unlike structure-based representation learning, which infers protein structures at various levels [32, 33, 34, 35, 36, 37], pLMs are capable of learning from the vastly larger corpus of sequenced proteins, which far exceeds those with experimentally validated structures [38, 39]. As pLMs scale in size, they have demonstrated the ability to produce robust protein representations for downstream prediction tasks that are competitive with leading methods like AlphaFold2 without the need for multiple-sequence alignments [26]. Variational Autoencoders (VAEs) have been utilized to infer phylogeny and generate libraries that enrich functional variants [15], while the CLEAN model employs contrastive learning with the InfoNCE loss to categorize enzymes based on their Enzyme Commission (EC) numbers, effectively predicting the reactions of uncharacterized sequences [40]. pLMs are not limited to particular protein families and can be trained over diverse protein types, including enzymes, binding domains, membrane receptors, and transporters [7] to learn transferable principles underpinning the syntax of protein sequences and offer great versatility in classifying and predicting diverse protein functions. Our work extends the use of pLMs by pretraining them to develop a joint latent embedding with a biomedical language model (bLM) that aligns protein sequences with natural language annotations to expose new understanding of protein homology and function and facilitate the guided design of proteins using text prompts.

### Multimodal Representation Learning

Leveraging natural language and textual representations has significantly enhanced representation learning across various modalities such as images [41, 42, 43, 44], video [45, 46, 47], speech [48, 49], and small molecules [50, 51]. Notably, recent advancements have aligned protein Language Models (pLMs) with biomedical Language Models (bLMs) using a multimodal InfoNCE loss originally implemented with the CLIP model [41] to attract and repel text-protein pairs in a joint embedding space [22, 23]. Herein, we build upon this prior work by increasing the training data abundance to 45M text-protein pairs curated from the SwissProt and Pfam databases [38], enriching the text prompts to incorporate additional protein attributes, and introducing a protein family contrastive loss to better infer homology within the joint embedding space.

### Conditional Generation for Protein Design

Generative protein design has historically been engaged through structure-based and sequence-based paradigms. In structure-based design, Denoising Diffusion Probabilistic Models (DDPMs) have emerged as powerful tools and have been applied to protein design [52] and secondary structure and fold engineering [53, 54]. RFDiffusion has been produced experimentally validated novel structures conditioned on desired functional motifs [55], and Chroma has shown success in conditional protein design using classifier guidance [56]. Sequence-based generation appeals to the sequence-structure-function paradigm wherein sequence encodes the salient information needed to predict both structure and function [1, 2, 3, 4, 5], and offers advantages in generative protein design relative to structure-based paradigms in the size of training data (𝒪 (10^9^) known sequences vs. 𝒪 (10^5^) solved structures) [38, 39], elimination of the need for inverse-folding models since sequences are directly designed [20, 57], and seamless integration with high-throughput functional enrichment assays using next-gen sequencing to probe sequence-function relationships [58]. Sequence-based generative models, such as autoregressive models [59, 60] and VAEs [15, 61], have shown experimental validation but are often limited by their conditioning capabilities. For example, ProGen uses control tags such as gene ontology and species labels to guide generation with an autoregressive language model [16], yet lacks the flexibility and simplicity of natural language prompting of conditional image generation models like DALL-E [43]. ProtST [22] aligns text embeddings in a joint space but has not been used for conditional generation with a protein decoder, whereas ProteinDT [23] generates *in silico* validated proteins using both autoregressive and discrete diffusion decoders, allowing for flexible sequence generation guided by text prompts. Our work leverages a richer and larger dataset of 45M text-protein pairs and implements an order-agnostic autoregressive diffusion model conditioned on textual descriptions that enables full sequences design or in-painting of desired regions similar to EvoDiff [24], but with the added capability of guided protein design through conditional generation using text prompts. Lastly, our work demonstrates the first demonstration, to the best of our knowledge, of the *in vivo* and *in vitro* validation of functional proteins designs based on natural language text prompts.

## 3 Methods

We summarize methodological details of curation of text-protein pairs, the three components – Protein embeddings with natural language using Contrastive Learning (PenCL), refinement and enhancement of the joint embedding (Facilitator), and generation of novel sequences via text-conditioned autoregressive discrete diffusion (ProteoScribe) – of the BioM3 model, and the *in vivo* and *in vitro* validation of designed SH3 sequences. Additional information is provided in Appendix A.

### 3.1 Dataset Curation and Assembly

We curated a corpus of 45M pairs of protein sequences and text annotations from the SwissProt and Pfam databases [38]. Similar to ProtST [22], we retrieved information from SwissProt and added prefixes to delimit annotations from different fields: “PROTEIN NAME”, “FUNCTION”, “SUBCELLULAR LOCATION”, and “SIMILARITY”. We also gathered information for additional prefixes, including: “CATALYTIC ACTIVITY”, “DOMAIN”, “LINEAGE”, “FAMILY NAME”, “ACTIVITY REGULATION”, “BIOPHYSICOCHEMICAL PROPERTIES”, “PTM”, “TISSUE SPECIFICITY”, “MISCELLANEOUS”, “COFACTOR”, “PATHWAY”, “BIOTECHNOLOGY”, and “INDUCTION”. We expanded our training data by appealing to the Pfam database, which sorts domains from sequences in UniProt and assembles protein families and clans. For these data, we added two more prefixes: “FAMILY DESCRIPTION” and “GENE ONTOLOGY”. Since the Pfam database is assembled based on protein families, the “FAMILY DESCRIPTION” field enriches our text prompt, although the previously mentioned fields might not always be available. Statistics on the availability of information for each field from the two databases is detailed in Tables S1 and S2. The full text description for each protein sequence is formed by concatenating all fields into a single prompt. Examples of text-protein pairs from the SwissProt and Pfam databases are presented in Tables S3 and S4. Additional details on the data curation are provided in Appendix A.1.

### 3.2 Stage 1: Protein embeddings with natural language using Contrastive Learning (PenCL)

During PenCL pretraining, we aim to infer a joint embedding space that aligns protein representations with natural language representations of textual protein descriptions. First, an intermediate protein representation *h*_*p*_ is inferred from a single-modality pretrained protein Language Model (pLM) and then further transformed into a protein representation *z*_*p*_ within the joint embedding space using a simple multi-layer perceptron module, often referred to as a projection head [62]. Similarly, we infer an intermediate text representation *h*_*t*_ using a single-modality pretrained biomedical Language Model (bLM) and then further transform it into a text representation *z*_*t*_ within the joint embedding space using a projection head. For the pLM, we used ESM2 [26] with 650M parameters, pretrained on protein sequences from the UniProt database with a masked language model loss. For the bLM, we used PubMedBERT-full [63] with 100M parameters, trained on PubMed full-text articles to extract representations of arbitrary textual descriptions.

To induce multimodal pretraining and align text representations with protein representations, we leverage the InfoNCE loss [64], originally implemented on multimodal text-image data with CLIP [41], and we call this objective, originally coined by ProtST [22], the Global Contrastive loss *L*_*GC*_:

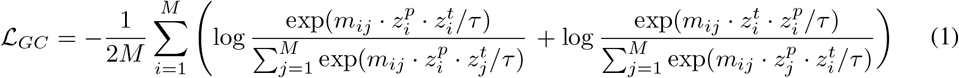

where *m*_*ij*_ is a masking matrix defined as:

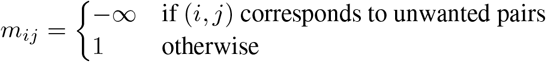

where *τ* is a temperature hyperparameter, *M* equals 2*N* where *N* is the batch size for SwissProt and Pfam and we curate the batch such that each sequence *i* = 1…*N* in the SwissProt batch corresponds to a sequence *i* = (*N* + 1)…*M* in the Pfam batch based on homology (i.e., matching Pfam labels), *z*_*t*_ is the text representation, and *z*_*p*_ is the protein representation. The masking matrix *m*_*ij*_ ensures that specific pairs do not contribute by setting the value to−∞, effectively excluding them from the calculation. This exclusion is necessary to eliminate false negatives corresponding, for example, to homologs *i* = *k* and *j* = *N* + *k* sampled from the SwissProt and Pfam databases. This masking aligns with the pretraining approach, avoiding misleading associations between particular pairs while still enforcing robust contrastive learning.

We also introduce a homology contrastive loss to take advantage of the homology information contained in the Pfam database that we term the Protein Family Contrastive Loss ℒ _*P F C*_:

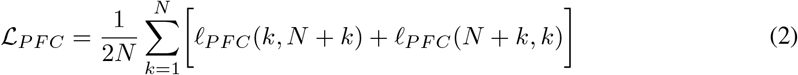

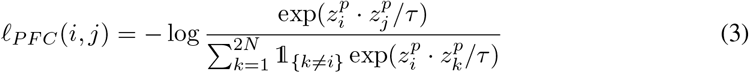

where *τ* is a temperature hyperparameter, *N* is the batch size, 𝟙 {_*k* ≠ *i*}_ is an indicator function, and *z*_*p*_ is the protein representation. This loss further attracts (repels) homologs from the same (different) protein families as classified by Pfam, thereby introducing a molecular evolution inductive bias. Along with these two contrastive losses, we continue to train the bLM and pLM with standard masked language model losses ℒ_*bML*_ and ℒ_*pML*_, further improving it’s ability to reconstruct protein sequences and our curated textual prompts. Thus, the total loss for PenCL is ℒ_*PenCL*_ = ℒ_*GC*_ + ℒ_*P F C*_ + ℒ_*bML*_ + ℒ_*pML*_, which we minimize with respect to the weights of the bLM, pLM, and the two projection heads. The model is trained over text-protein batches curated from SwissProt and Pfam and infers them in parallel with the bLM and pLM (Figs. S2, S3, S6A). More details on the architecture and lossess are provided in Appendix A.2.1, pretraining details are provided in Appendix A.2.2, and ablation analysis details are provided in Appendix A.2.3.

### 3.3 Stage 2: Facilitator

The second stage is to use a module referred to as a Facilitator, which acts as an alignment module on top of PenCL to further improve the text embeddings. This module was originally introduced by ProteinDT [23] and was found to empirically improve sequence generation. The Facilitator is a simple autoencoder architecture, where the input is the text embedding *z*_*t*_, the output embedding is an augmented text embedding *z*_*c*_, and the objective is to reconstruct *z*_*p*_. Instead of a mean-square error loss similar to ProteinDT, we used a max-mean discrepancy (MMD) loss to compute the similarity between the distributions of *z*_*p*_ and *z*_*c*_. MMD metric operates in a high-dimensional feature space, enabling more nuanced comparisons and better handling of high-dimensional data [65]. The Stage 2 training loss objective for the Facilitator is given by:

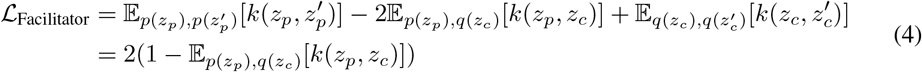

where *p*(*z*_*p*_) is the distribution over *z*(_*p*_, *q*(*z*_*c*_) is the distribution over *z*_*c*_, and *k*(·, ·) is a Gaussian kernel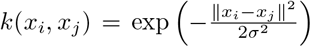 where *σ* is a hyperparameter. The simplification after the second equality results from the fact that the first and third terms evaluate to unity. We minimize the loss function with respect to the weights of the Facilitator model. Alternatively, we can replace MMD loss and compute the mean-squared error (MSE) loss between *z*_*p*_ and *z*_*c*_, although the MMD loss appears to be more robust and less prone to overfitting (Fig. S4A). Empirically, we found that a key function of the Facilitator is to improve the agreement between the norm of text embedding || *z*_*t*_|| and protein embedding || *z*_*p*_ || (Fig. S4B). More details of the architecture and loss objectives are provided in Appendix A.3.1, and pretraining details are provided in Appendix A.3.2.

### 3.4 Stage 3: Protein Generation with Textual Descriptions (ProteoScribe)

To generate artificial sequences that are compatible with a given text prompt, we pretrain a decoder model on the text-protein pairs from the SwissProt database. While ProteinDT [23] has shown that sequences can be generated by conditioning on a text prompt with an autoregressive language decoder or discrete diffusion decoder, we instead choose to implement a discrete order-agnostic autoregressive diffusion model (ARDM) [66]. This architecture allows us to flexibly generate complete sequences or in-paint arbitrary (non-contiguous) motifs in an order agnostic fashion. EvoDiff [24] presented an early implementation of ARDM for proteins, and we expand upon this architecture by incorporating conditional capabilities to enable the model to accept conditioning on natural language text prompts. The pretraining process involves sampling time points *t*, corrupting protein sequences *x* with absorbing states based on these sampled time points, and denoising the remaining sequence by reconstruction while conditioned on both the uncorrupted sequence tokens and the *z*_*c*_ embedding (Figure S5). Thus, loss objective for ProteoScribe is given by:

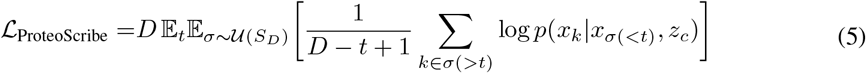

where *D* is the number of timesteps, *t* is the timestep, 𝒰 (*S*_*D*_) is the uniform distribution of all possible orderings of positions *σ* ∈ *S*_*D*_, *x* is the protein sequence, *x*_*k*_ is the *k*^th^ token of the protein sequence, *x*_*σ*(*<t*)_ is the set of (possibly non-contiguous) tokens specified or decoded prior to time *t*, and *z*_*c*_ = *f*_Facilitator_(*z*_*t*_) is the protein sequence representation produced by the Facilitator. We implement this as an efficient transformer model [67, 68, 69]. Further details of the architecture and loss objectives are provided in Appendix A.4.1, pretraining details are provided in Appendix A.4.2, and the sampling procedures for generating sequences or in-painting motifs is provided in Appendix A.4.3.

### 3.5 Experimental Validation of Prompt Engineering for Proteins

We experimentally validate the trained BioM3 model by conducting *in vivo* and *in vitro* experimental tests of generated protein sequences conditioned on text prompts containing information about Src homology 3 (SH3) sequences capable of functioning like natural Sho1^SH3^ domains by binding their cognate pbs2 ligand and effecting the osmosensing mechanism in *S. cerevisiae* (baker’s yeast). To do so we fine tuned the ProteoScribe model on a curated library of 25,030 SH3 domain-containing sequences. The fine-tuned model thus integrates the text-protein alignment capabilities of the PenCL and Facilitator stages with the specialized knowledge of SH3 domain structure and function within ProteoScribe. After fine-tuning is complete, we constructed five text prompts of varying similarity to SwissProt and Pfam annotation styles and deployed the BioM3 model to generate artificial sequences. The generated sequences – 984 across all five prompts – were then subjected to gene synthesis, assembly, and *in vivo* and *in vitro* tests of function. We tested the functional capacity of the designed sequences to rescue activity and promote survival under selective conditions *in vivo* using a high-throughput select-seq assay that couples a high-osmolarity challenge with next-generation sequencing to measure the relative enrichment (r.e.) of the post-selection population in a particular allele variant relative to a null gene and wild-type *S. cerevisiae* [15]. The r.e. score provides a quantitative measurement of the degree to which our designed SH3 domains are functional *in vivo* and capable of activating a homeostatic osmoprotective response. Additionally, we selected a handful of designed sequences for purification and measurement of *in vitro* activity in a biochemical binding assay. Full details of the preparation of the SH3 dataset is presented in Appendix A.5.1, the fine-tuning of ProteoScribe in Appendix A.5.2, the five text prompts and sampling of designed sequences in Appendix A.5.3, and the experimental details of the *in vivo* and *in vitro* assays in Appendix A.5.4 and Appendix A.5.5.

## 4 Experiments and Results

We conducted a series of tests to evaluate the performance of our BioM3 framework in designing functional protein sequences guided by natural language prompts. Our results demonstrate the efficacy of our approach across multiple dimensions, including *in silico* validations of subcellular localization prediction, reaction classification, remote homology detection, scaffold in-painting, and structural plausibility, and *in vivo* and *in vitro* assays of natural language prompt-designed synthetic analogs of Src-homology 3 (SH3) domain proteins.

### 4.1 Visualization of PenCL Protein-Text Embedding

The PenCL architecture effectively integrates protein sequences and text descriptions into a unified joint embedding space. The Global Contrastive Loss (ℒ_*GC*_) aligned text and protein embeddings, while the Protein Family Contrastive Loss (ℒ _*P F C*_) ensured that homologous sequences from the same family were closely clustered (Fig. S6A). The clustering performance was assessed using the Calinski-Harabasz Index (CHI) and Davies-Bouldin Index (DBI) [70], with results indicating higher clustering quality for PenCL compared to single modality models (Fig. S6B). A Principal Component Analysis (PCA) embedding of the joint embedding space exposed distinct clustering of protein families based on their textual descriptions and a clear separation of inter-clan and intra-clan clusters (Fig. S7), thereby confirming the model’s ability to infer homology and functional relationships from the text-protein pairs.

### 4.2 Benchmarking PenCL in Zero-Shot and Homolog Detection Tasks

PenCL’s performance was further benchmarked in zero-shot and homolog detection tasks, focusing on subcellular localization prediction, reaction classification, and remote homology detection. In zero-shot evaluations, PenCL demonstrated excellent performance in predicting subcellular localization and enzyme reaction classification, achieving accuracies on par with or exceeding that of the ProST [22] multimodal model (Table S5). For zero-shot remote homology detection, PenCL outperformed BLASTp [71], ProtST [22], ProtT5 [29], and ESM2 [29] in identifying homologous relationships between proteins, highlighting the performance capabilities realized by leveraging joint text-sequence embeddings for evolutionary and functional inference (Table S6). In addition, an ablation analysis was conducted on the remote homology detection task, demonstrating that the ℒ_*P F C*_ plays a crucial role in performance (Table S7). These benchmarks underscore PenCL’s robustness and versatility in handling diverse and complex protein-related tasks without fine-tuning. More details on the results and capabilities of PenCL are presented in Appendix B.1.

### 4.3 Sequence Generation and Structural Plausibility

We next tested the capability of the trained BioM3 model to produce structurally plausible protein sequences across a wide range of lengths and complexities based on distinct text prompts. In Figure S8, we present five text prompts drawn from the SwissProt training data and passed to BioM3 as an in-distribution test of its capacity to recapitulate protein sequence and structure within the training set. Structures were predicted *in silico* using ColabFold [72] and compared with experimentally solved or predicted structures retrieved from the AlphaFold2 database [73] corresponding to the text prompt. TMScores exceeding 0.922 and RMSD values below 2.32 Å indicate excellent agreement between the structure of the native sequence and that produced by BioM3. We also retrieved the nearest sequence using BLASTp to demonstrate that in all cases we recover a sequence with the same name and function as contained within the text prompt, but that the generated sequences possess high sequence diversity, sharing as little as 53.59% sequence similarity with the nearest BLASTp hit. We also tested the capacity of BioM3 for scaffold in-painting by retaining the functional motif of a compact calmodulin protein and generating novel scaffold regions conditioned on the sequence of the functional motif and a calmodulin text prompt (Appendix B.3). The functional motif remains consistent across all designs while generating diverse scaffoldings. These results showcase the expressive capabilities of text-guided generation, underscoring the potential in flexible and precise protein design. More details on the capabilities of ProteoScribe are presented in Appendix B.3.

### 4.4 Experimental Validation of Text-Guided Protein Designs

As a final test, we evaluated the *in vivo* and *in vitro* functionality of SH3 domain sequences generated using five distinct text prompts (Fig. 2A). The five prompts are presented in Appendix A.5.3. The first corresponds to the prompt annotation for Sho1^SH3^ that was seen during fine-tuning. The second and third are ablated versions of the first: the second prompt retains functional information, whereas the third includes only the name of the Sho1^SH3^ domain. The fourth and fifth prompts were entirely new: the fourth includes only the protein name, and the fifth incorporates the name and functional description. The trained BioM3 model with a ProteoScribe block fine-tuned on SH3 was primed with each of these prompts and generated 200, 200, 67, 200, and 317 artificial sequences with prompt 1, 2, 3, 4, and 5, with 984 prompt designs selected for experimental validation (Section 3.5, Appendix A.5.3). These sequences were then subjected to *in vivo* tests of osmosensing function by assessing the capacity of *S. cerevisiae* cells containing these designed protein variants to survive under high-osmolarity conditions as quantified by the relative enrichment (r.e.) scores measured in a high-throughput select-seq assay (Fig. 2B, Appendix A.5.4). The sequences generated by the various prompts exhibited a range of sequence identities compared to the closest natural SH3 domain spanning approximately 20-80%. Interestingly, the sequence identities are quite tightly clustered for each prompt, but the r.e. scores exhibit a range of values. Notably, sequences generated from prompts that included both name and functional descriptions tended to produce more functional alleles (Prompts 1, 2, and 5), while less descriptive prompts resulted in more novel sequences (Prompts 3 and 4) achieving as little as 40% identity while preserving residual function (Fig. 2C). We conducted low-throughput *in vitro* biochemical measurements of the binding affinity to the pbs2 ligand for four selected sequences from Prompt 5 spanning a range of r.e. values (Table S8). Notably, one design exhibited a significantly lower dissociation constant (i.e., higher binding affinity) of *K*_*d*_ = (0.31*±* 0.04) compared to 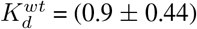 for the wild-type. Additionally, we observe that one functional design from Prompt 4 emerged as the most novel sequence to survive the selection media, having only two BLAST hits in the NCBI database (web browser) [74], neither of which returned a Sho1^SH3^ domain. Indeed, its closest identity to the wild-type Sho1^SH3^ was just 28% and to any natural SH3 domain in the library just 38% (Table S9). Further experimental statistics, data analysis, and discussion are provided in Appendix B.4. These results provide experimental confirmation that natural language prompts can effectively guide the design of unique and functional proteins.

**Figure 2:**
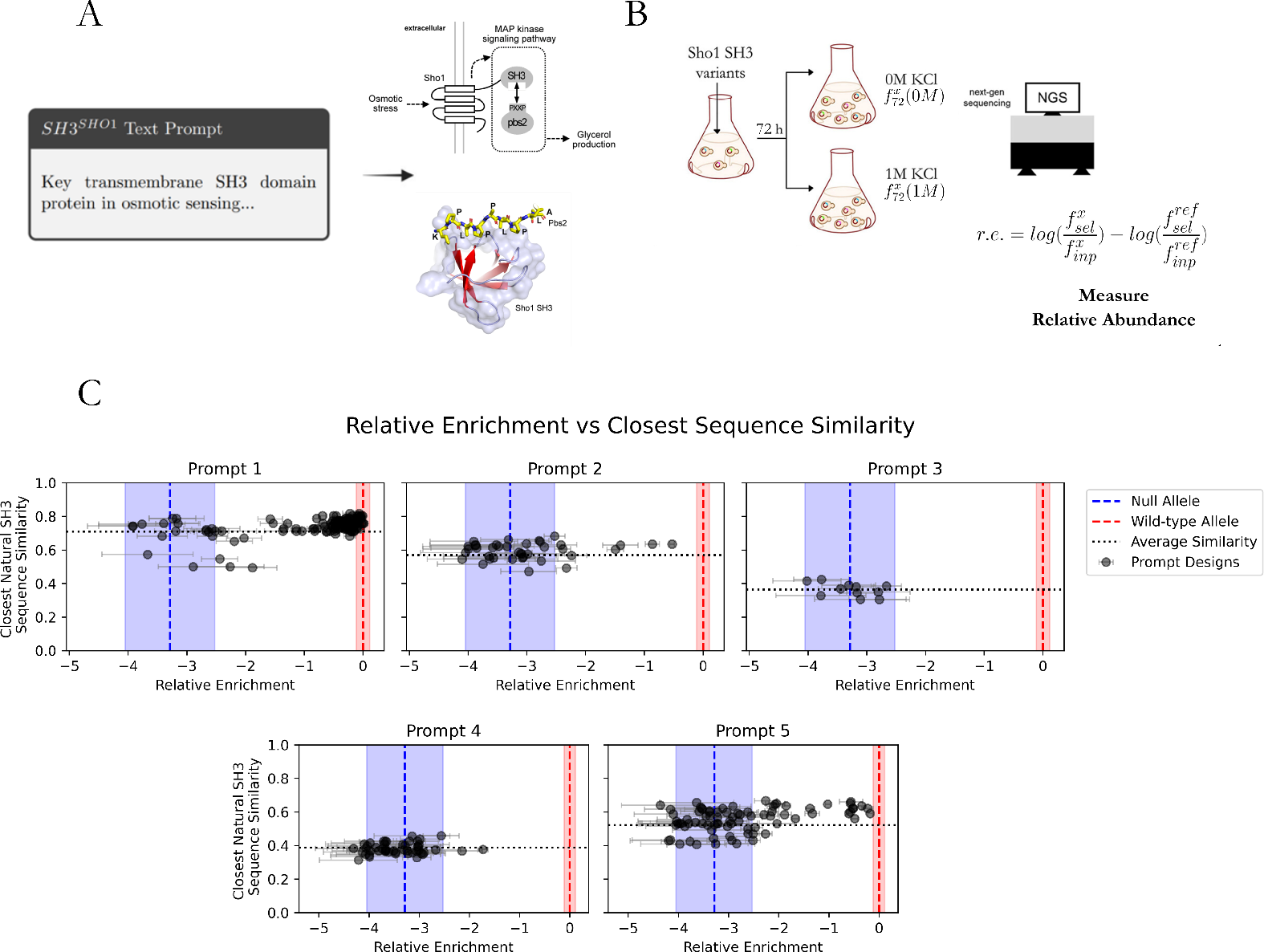
Experimental validation of BioM3 text-guided protein designs. (A) Illustration of text-guided protein design where the BioM3 model is prompted with a text description of the Sho1^SH3^ binding domain. SH3 domains are protein interaction modules that bind to polyproline-containing target ligands. For the Sho1 paralog, binding between the protein and the target peptide sequence in the pbs2 MAP kinase mediates responses to fluctuations in external osmotic pressure by controlling the production of internal osmolytes. We illustrate the experimentally solved structure of the *S. cerevisiae* Sho1^SH3^ domain (PDB:2VKN) in complex with the pbs2 peptide ligand. (B) Experimental assessment of the text-prompted, generatively-designed synthetic Sho1^SH3^ domains is conducted using a high-throughput select-seq assay that quantifies *in vivo* rescue of osmosensing function via the relative enrichment (r.e.) score. (C) Plots reporting the r.e. scores for the 984 sequences designed by the BioM3 model under each of the five prompts. The relative enrichment (r.e.) scores quantify the fitness of the designed proteins relative to wild-type (r.e. = 0) and the Levenshtein distance relative to the closest natural SH3 measures sequence divergence from natural. The blue dotted vertical line represents the r.e. = (−3.29 ±0.76) for a null allele, with the blue-highlighted interval indicating the uncertainty determined by error propagation. The red dotted vertical line shows the r.e. = (0.00± 0.11) corresponding to wild-type *S. cerevisiae* Sho1^SH3^, with its associated uncertainty also determined by error propagation highlighted in red. The various prompts induce different distributions of sequence divergence and osmosensing rescue. Notably, Prompt 1 and 5 both generate a number of sequences with wild-type-level function and as little as 47% sequence similarity. [Illustrations in A and B are adapted from Refs. [15] and [17].]

## 5 Conclusion and Future Work

In this work, we introduced BioM3 as a framework for the generative design of functional protein sequences conditioned on natural language prompts. By learning a joint embedding space between a protein language model (pLM) and a biomedical language model (bLM) and coupling this embedding as a conditioning to an autoregressive diffusion model, our approach effectively learns a mapping between protein sequence and text annotations and leverages this to guide the generation of novel sequences with desired functional attributes. Our *in silico* validations demonstrate the capability of the model to compete with or outperform state-of-the-art models for zero-shot prediction tasks and homology detection. We also present the first, to the best of our knowledge, *in vivo* and *in vitro* experimental tests of artificial proteins designed using natural language text prompts. In an application to the design of synthetic analogs of Src-homology 3 (SH3) domain proteins, all five prompts studied generated sequences with measurable function and revealed that prompts incorporating both name and functional descriptions tended to yield more functional designs. Notably, the very simple Prompt 5 that is quite distinct from the formal annotations in the training data – “Key transmembrane SH3 domain protein in osmotic sensing for filamentous growth and HOG pathways, involving Cdc42p/MAP kinase interactions and phosphorylation” – produced 25/317 sequences with comparable *in vivo* function to the wild-type Sho1^SH3^. In future work, we plan to expand our experimental tests of text-guided proteins to a wider class of protein types, including catalytic enzymes, transporter proteins, and therapeutic proteins, and perform scaffold in-painting to enable precise and flexible design of multi-domain and multi-functional proteins. The promising results of BioM3 underscore the powerful potential of combining natural language processing with protein design, offering a powerful tool for the intuitive and flexible creation of novel proteins and democratization of protein design tools to non-expert users through the use of simple natural language text prompts.

## Acknowledgements

We gratefully acknowledge support from the Machine Learning in the Chemical Sciences and Engineering program of The Camille and Henry Dreyfus Foundation (A.L.F.) and grant NIH RO1GM141697 from the National Institutes of General Medical Sciences (R.R.). This material is based upon the work supported by the National Science Foundation Graduate Research Fellowship Program under Grant No. 2140001 (N.P.). Any opinions, findings, and conclusions or recommendations expressed in this material are those of the author(s) and do not necessarily reflect the views of the National Science Foundation. We gratefully acknowledge the Medical Scientist Training Program (MSTP) Training Grant T32GM150375 (H.Y.). This work was completed in part with resources provided by the University of Chicago Research Computing Center. We gratefully acknowledge computing time on the University of Chicago high-performance GPU-based cyberinfrastructure supported by the National Science Foundation under Grant No. DMR-1828629. This research used resources of the Argonne Leadership Computing Facility, a U.S. Department of Energy (DOE) Office of Science user facility at Argonne National Laboratory and is based on research supported by the U.S. DOE Office of Science-Advanced Scientific Computing Research Program, under Contract No. DE-AC02-06CH11357.

## Appendix and Supplementary Materials

### A Detailed Methods

#### A.1 Data Curation, Assembly, and Statistics

To curate the pretraining dataset for BioM3, we integrated sequences and annotations from the SwissProt and Pfam databases [38]. This approach aimed to create a richly annotated dataset that could effectively guide text-to-protein generative tasks, allowing for the development of functionally relevant and structurally plausible protein sequences.

##### SwissProt Database Curation

We retrieved 569,516 sequences from SwissProt [38] and systematically annotated each with detailed textual fields. Standard fields utilized in previous models like ProtST included “PROTEIN NAME,” “FUNCTION,” “SUBCELLULAR LOCATION,” and “SIMILARITY.” To enhance the dataset, we introduced several additional fields such as “CATALYTIC ACTIVITY,” “DOMAIN,” “LINEAGE,” “FAMILY NAME,” “ACTIVITY REGULATION,” “BIO-PHYSICOCHEMICAL PROPERTIES,” “PTM,” “TISSUE SPECIFICITY,” “MISCELLANEOUS,” “COFACTOR,” “PATHWAY,” “BIOTECHNOLOGY,” and “INDUCTION.” These novel fields, highlighted in blue in Table S1, were not previously utilized in similar datasets, enrich the text prompts with comprehensive information about protein functions and properties.

##### Pfam Database Curation

We further expanded our dataset by incorporating the Pfam database [38], which organizes proteins into families and clans based on sequence homology and domain structure. The Pfam dataset included over 44 million sequences annotated with unique fields, such as “FAMILY DESCRIPTION” and “GENE ONTOLOGY,” highlighted in red in Table S2. These fields provide an added layer of biological context, describing family-specific functions and gene-level annotations. This integration of detailed family descriptions and gene ontology significantly broadens the dataset’s corpus, providing diverse and informative text prompts that are critical for effective model training.

##### Visualizing Database Content

To emphasize the differences in annotation content between SwissProt and Pfam, we generated word clouds for each database (Figure S1). The word clouds illustrate the prevalence of various keywords, with larger words indicating higher frequency. For SwissProt, terms like “ribosomal,” “cytoplasmic,” and “subunit” dominate, highlighting the database’s emphasis on subcellular localization and protein complexes. In contrast, the Pfam word cloud shows a broader array of terms, such as “binding,” “domain,” and “activity,” reflecting the detailed functional and structural annotations provided by the Pfam database. This visualization highlights how the combined use of SwissProt and Pfam data enriches the dataset, contributing to the model’s ability to generate functionally and structurally diverse proteins. The distinct word clouds underscore the complementary nature of these databases in providing a comprehensive set of annotations for protein sequence generation.

##### Statistics and Overview

Tables S1 and S2 summarize the statistics of the SwissProt and Pfam databases, respectively, highlighting the distribution of different annotation fields. The SwissProt dataset shows a high prevalence of functional descriptions, with 81% of sequences having a defined function, while the Pfam dataset is characterized by detailed family annotations, with 100% of sequences labeled with “FAMILY DESCRIPTION.” The unique keywords retrieved from both databases, particularly those new to our dataset, play a crucial role in enhancing the richness and diversity of text prompts for pretraining.

##### Example Text-Protein Pairs

Tables S3 and S4 provide examples of text-protein pairs from SwissProt and Pfam, demonstrating the dataset’s ability to capture comprehensive biological information. Each pair includes a protein sequence aligned with a corresponding text description generated by concatenating all available fields, showcasing the breadth of annotations and their informational capacity to guide generative models in creating biologically relevant and functional protein sequences.

**Figure S1:**
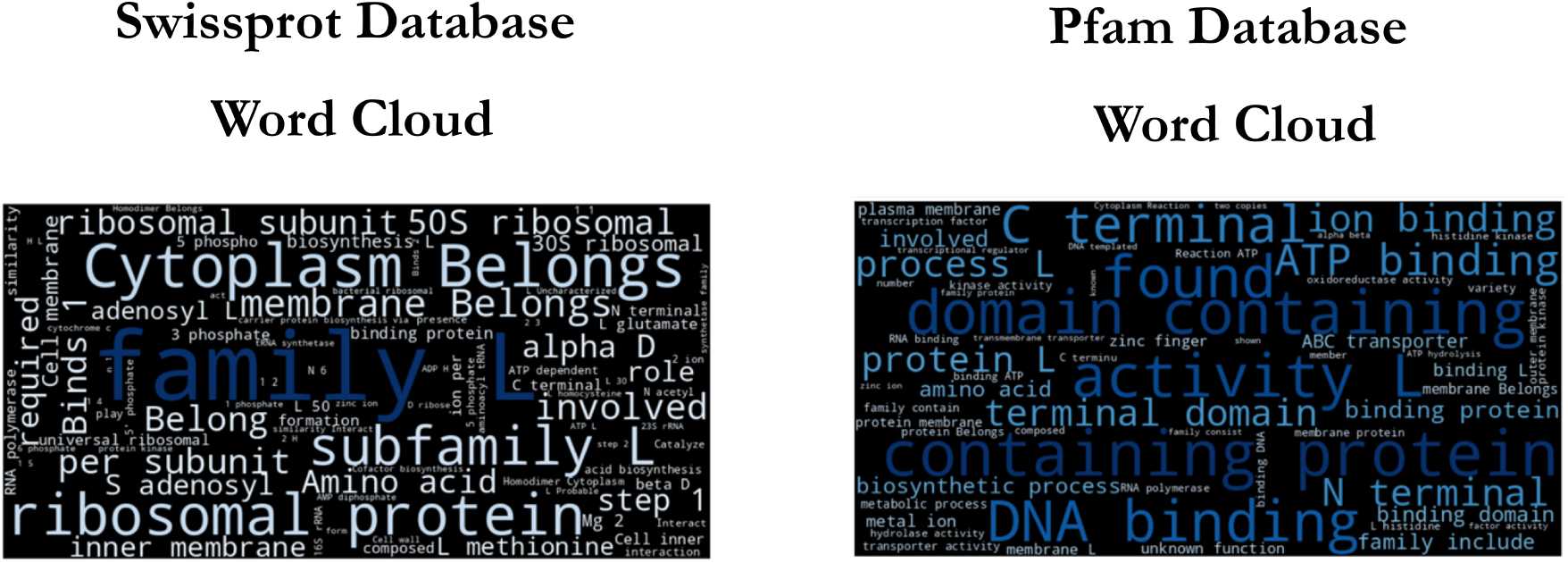
Word clouds visualizing the most frequent terms in the SwissProt (left) and Pfam (right) databases [38]. The size of each word corresponds to its frequency in the respective database. The SwissProt word cloud emphasizes terms related to cellular components and protein classification, with prominent words like “Cytoplasm,” “Belongs,” and “family.” The Pfam word cloud highlights functional and structural terms, with “domain,” “binding,” and “activity” being notably prominent. These visualizations illustrate the complementary nature of the two databases in providing comprehensive protein annotations for the BioM3 training dataset.

**Table S1:**
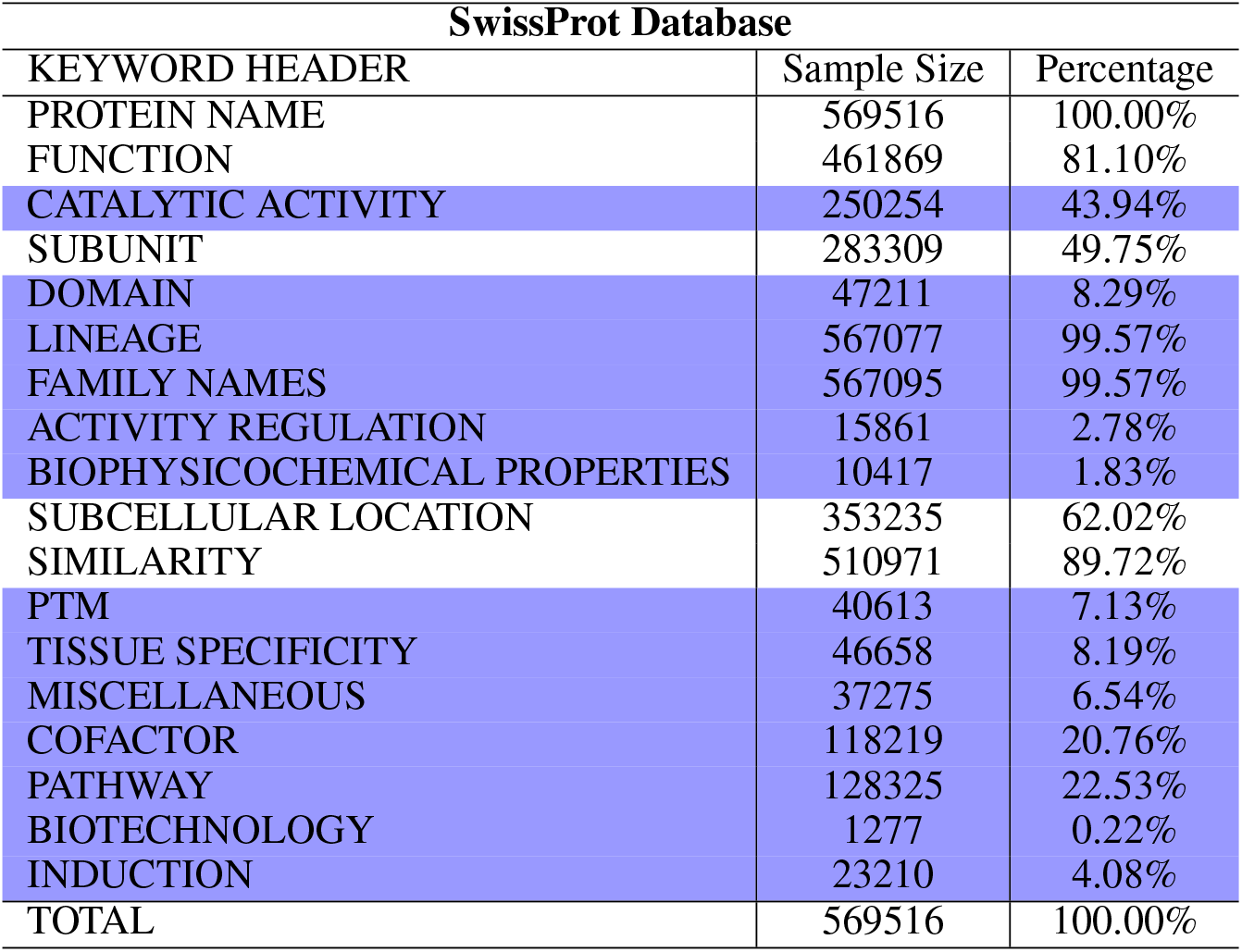
Distribution of keyword headers in the SwissProt Database, showing sample sizes and percentages for various annotation fields. Blue highlighted keywords represent novel information added to text prompts compared to datasets curated by ProtST [23] and ProteinDT [22].

| KEYWORD HEADER | Sample Size | Percentage |
| --- | --- | --- |
| PROTEIN NAME | 569516 | 100.00% |
| FUNCTION | 461869 | 81.10% |
| CATALYTIC ACTIVITY | 250254 | 43.94% |
| SUBUNIT | 283309 | 49.75% |
| DOMAIN | 47211 | 8.29% |
| LINEAGE | 567077 | 99.57% |
| FAMILY NAMES | 567095 | 99.57% |
| ACTIVITY REGULATION | 15861 | 2.78% |
| BIOPHYSICOCHEMICAL PROPERTIES | 10417 | 1.83% |
| SUBCELLULAR LOCATION | 353235 | 62.02% |
| SIMILARITY | 510971 | 89.72% |
| PTM | 40613 | 7.13% |
| TISSUE SPECIFICITY | 46658 | 8.19% |
| MISCELLANEOUS | 37275 | 6.54% |
| COFACTOR | 118219 | 20.76% |
| PATHWAY | 128325 | 22.53% |
| BIOTECHNOLOGY | 1277 | 0.22% |
| INDUCTION | 23210 | 4.08% |
| TOTAL | 569516 | 100.00% |

**Table S2:** Summary statistics of Pfam database entries, showing sample sizes and percentages for various keyword headers. Blue highlighted keywords represent novel information added to text prompts compared to datasets curated by ProtST [23] and ProteinDT [22]. Red highlighted keywords are uniquely retrieved from the Pfam database, allowing expansion of the text-protein dataset corpus while maintaining meaningful text prompts.

| Pfam Database |  |  |
| --- | --- | --- |
| KEYWORD HEADER | Sample Size | Percentage |
| PROTEIN NAME | 42797279 | 95.56% |
| FUNCTION | 6426970 | 14.36% |
| CATALYTIC ACTIVITY | 5893975 | 13.16% |
| SUBUNIT | 3277681 | 7.32% |
| DOMAIN | 458266 | 1.02% |
| LINEAGE | 43088859 | 96.25% |
| FAMILY NAMES | 44767154 | 100.00% |
| ACTIVITY REGULATION | 81415 | 0.18% |
| BIOPHYSICOCHEMICAL PROPERTIES | 15681 | 0.03% |
| SUBCELLULAR LOCATION | 9418016 | 21.04% |
| SIMILARITY | 17398665 | 38.86% |
| PTM | 190700 | 0.04% |
| TISSUE SPECIFICITY | 0 | 0.00% |
| MISCELLANEOUS | 179241 | 0.40% |
| COFACTOR | 4358989 | 9.74% |
| PATHWAY | 3089004 | 6.90% |
| BIOTECHNOLOGY | 0 | 0.00% |
| INDUCTION | 16136 | 0.04% |
| FAMILY DESCRIPTION | 44767154 | 100.00% |
| GENE ONTOLOGY | 43088859 | 96.25% |
| TOTAL | 44767154 | 100.00% |

**Table S3:**
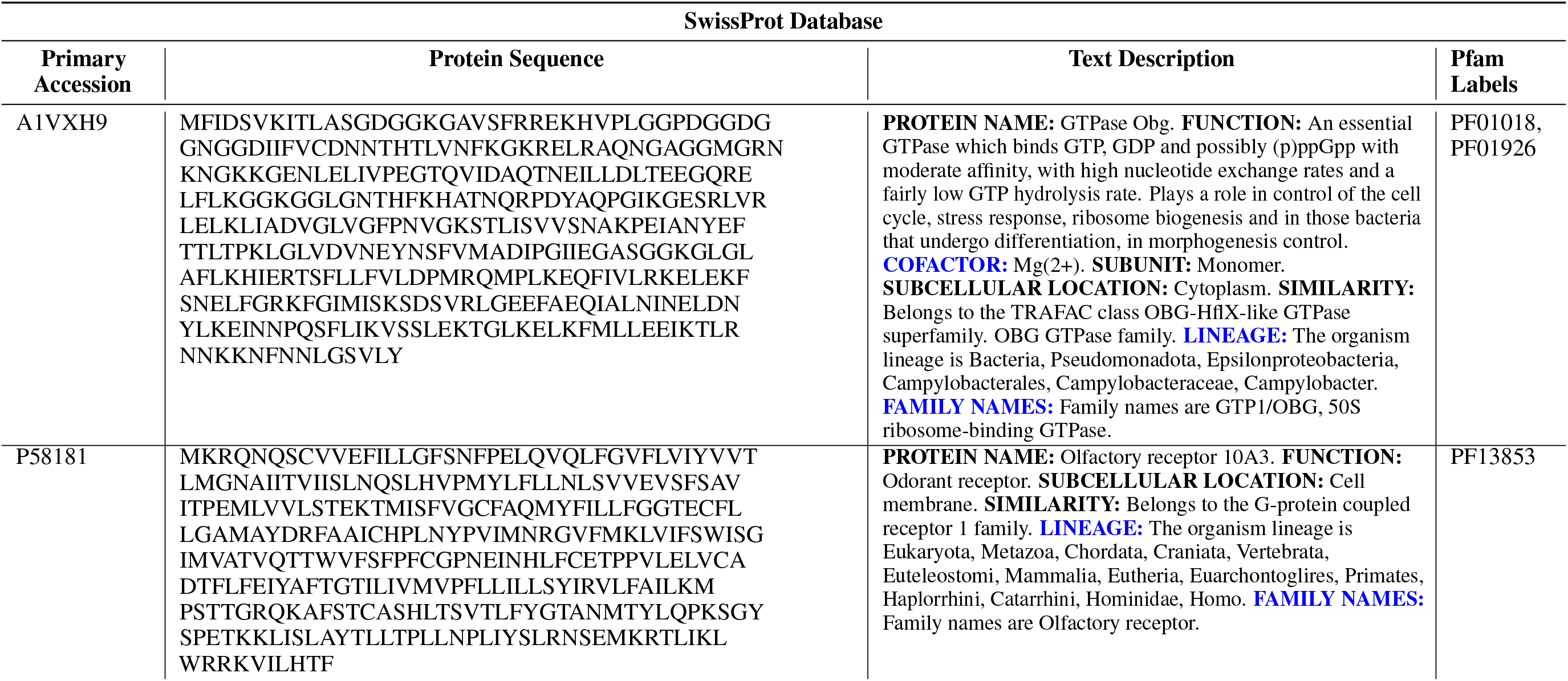
Examples of text-protein pairs from the SwissProt database [38]. This table illustrates the alignment of protein sequences with their corresponding detailed text descriptions. The blue text in the Text Description column indicates headers that were uniquely retrieved for our database, expanding the annotation beyond standard fields. The table includes various annotation fields such as protein name, function, subcellular location, and family names. The rightmost column shows associated Pfam labels.

**Table S4:**
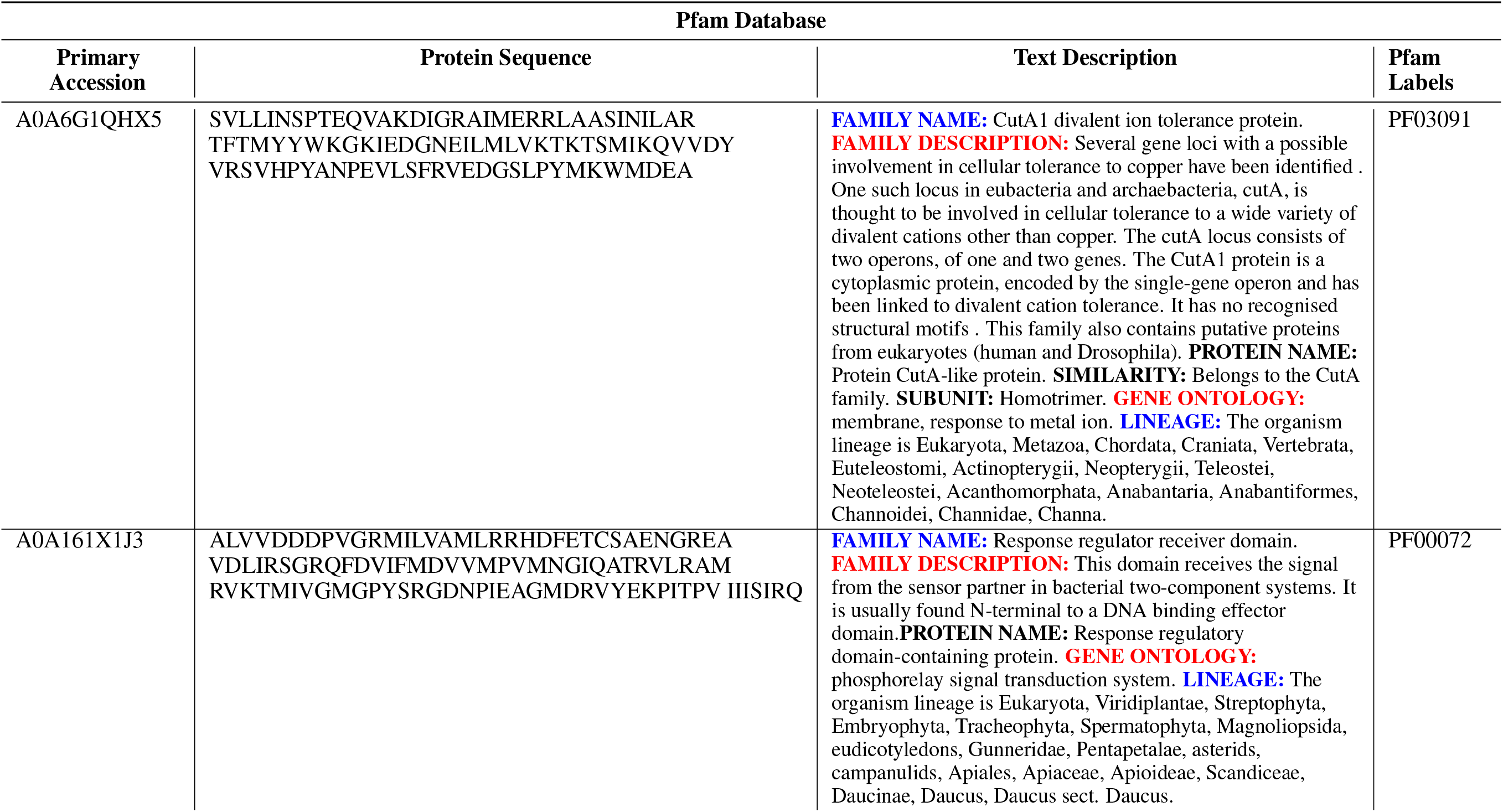
Examples of text-protein pairs from the Pfam database [38]. This table demonstrates the alignment of protein sequences with their corresponding detailed text descriptions. The red text in the Text Description column highlights headers uniquely retrieved from the Pfam database, such as “FAMILY DESCRIPTION” and “GENE ONTOLOGY”, which expand the annotation corpus. Blue text indicates other important annotation fields. The table includes various annotation fields such as family name, protein name, similarity, and lineage. The rightmost column shows the associated Pfam labels.

#### A.2 Stage 1: Protein embeddings with natural language using Contrastive Learning (PenCL)

##### A.2.1 Architecture and Loss of PenCL

The first stage of our model framework creates a joint embedding space aligning protein representations with natural language representations of textual protein descriptions. This stage, termed PenCL (Protein embeddings with natural language using Contrastive Learning), forms the foundation of our model’s ability to understand and align protein sequences with their textual descriptions.

Our approach employs two separate language models: a protein Language Model (pLM) and a biomedical Language Model (bLM). The pLM is based on ESM2 [26] with 650M parameters, while the bLM utilizes PubMedBERT-full [63] with 100M parameters.

The embedding process occurs in two steps for each modality. For proteins, an intermediate protein representation (*h*_*p*_) is first inferred from the pLM. For text, an intermediate text representation (*h*_*t*_) is inferred from the bLM. Importantly, both *h*_*p*_ and *h*_*t*_ are taken as the first hidden token from their respective language models. These representations are then transformed into a protein representation (*z*_*p*_) and a text representation (*z*_*t*_) within the joint embedding space using multi-layer perceptron modules, referred to as projection heads [62]. Thus the inference to acquire representations for the protein sequences has the following pipeline:

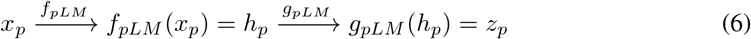

where *x*_*p*_ is the protein sequence input features, *f*_*pLM*_ is the functional mapping from the protein Language Model (pLM), *h*_*p*_ is the hidden representation along the starting token position that the transformer-based pLM infers, *g*_*pLM*_ is the projection head, and *z*_*p*_ is the final latent protein sequence representation in the joint embedding space. Similarly, for the inference to acquire representations for text prompts has the following pipeline:

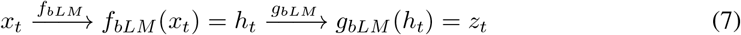

where *x*_*t*_ is the protein sequence input features, *f*_*bLM*_ is the functional mapping from the biomedical Language Model (bLM), *h*_*t*_ is the hidden representation along the starting token position that the transformer-based bLM infers, *g*_*bLM*_ is the projection head, and *z*_*t*_ is the final latent text representation in the joint embedding space.

The overall loss function for our PenCL architecture is defined as follows:

###### Stage 1 Pretraining Loss

The Stage 1 pretraining loss objective for PenCL is the following:

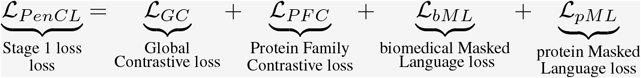

where

ℒ _*GC*_ → Learns representations that align protein sequence and natural text.

ℒ → _*P F C*_ Learns representations that are invariant to various homologs from the same family, while also being able to distinguish between different families.

ℒ → _*bML*_ + ℒ _*pML*_→ Learns to infer a masked word (amino acid) from its context, learning rich language representations that encompass word (amino acid) meaning and grammatical (coevolution) relationships.

The components of this loss function are defined as follows: Stage

###### The Stage 1 pretraining loss equations components

1 Pretraining Loss Equation Components

The Stage 1 pretraining loss objective for PenCL is defined by the following components:

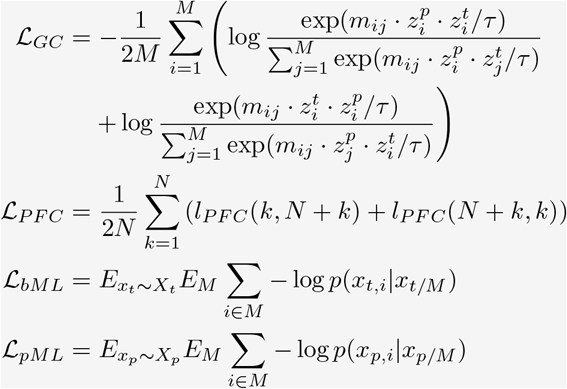

where

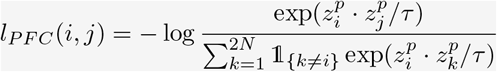

*τ* is a temperature parameter that scales the distribution of the dot products, and *m*_*ij*_ is a masking matrix defined as:

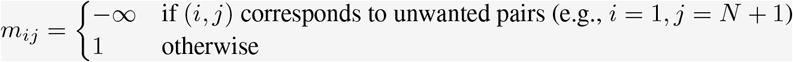

𝟙 {_*k≠i*_} is the indicator function ensuring that masked entries do not contribute as negative pairs.

In these equations, *τ* is a temperature hyperparameter, *M* = 2*N* where *N* is the batch size for SwissProt and Pfam, *z*_*t*_ is the text representation, and *z*_*p*_ is the protein representation. Importantly for implementation of our loss functions, we curate the batch such that each sequence *i* = 1…*N* in the SwissProt batch corresponds to a sequence *i* = (*N* + 1)…*M* in the Pfam batch based on homology (i.e., matching Pfam labels) (cf. Supplementary Algorithm 1). The Global Contrastive loss (ℒ_*GC*_) aligns text representations with protein representations. The Protein Family Contrastive Loss (ℒ_*P F C*_) leverages homology and the structure of the Pfam database, introducing a molecular evolution inductive bias into our model architecture. The masked language model losses (ℒ_*bML*_ and ℒ_*pML*_) for both the bLM and pLM allow our model to simultaneously optimize for protein sequence reconstruction and textual prompt understanding.

This comprehensive loss function enables our model to create a unified representation space where proteins and their textual descriptions can be meaningfully compared and analyzed, setting the stage for various downstream tasks in our overall framework.

##### A.2.2 PenCL Pretraining

The PenCL model is pretrained to align protein sequences with their corresponding textual descriptions using a contrastive learning framework that integrates ESM2, a protein language model, and PubMedBERT, a biomedical language model. This process forms a joint embedding space that not only captures the complex relationships between proteins and their natural language descriptions but also contrasts homology to leverage molecular evolution inductive bias. This dual approach is especially important given the absence of an exhaustive large-scale protein-text data corpus, allowing PenCL to effectively generalize across protein-text modalities and utilize evolutionary context for tasks such as functional annotation and protein design.

Figure S2 provides an overview of PenCL’s joint embedding and masked language loss objectives. The figure illustrates how protein sequences from the Swiss-Prot and Pfam databases are processed to compute both the Global Contrastive loss (ℒ_*GC*_) and Protein Family Contrastive loss (ℒ_*P F C*_). For each training pass, a batch size *N* is sampled from Swiss-Prot, and sequences from Pfam with matching Pfam labels are selected to form positive and negative pairs. This setup allows the simultaneous computation of both ℒ_*GC*_ and ℒ_*P F C*_ in a single forward pass, efficiently leveraging labeled data to train PenCL. The masked language modeling losses (ℒ_*bML*_ and ℒ_*pML*_) are computed using sequences with masked tokens, enhancing the model’s ability to capture contextual and semantic relationships within both the protein sequences and textual descriptions.

The dataloader algorithm (Supplementary Algorithm 1) details the sampling process of protein-text pairs along with their homologs, while the pretraining algorithm (Supplementary Algorithm 2) describes the step-by-step workflow involved in PenCL’s training. As detailed in the algorithm, each minibatch of protein and text data is processed through the pLM and bLM to generate intermediate representations (*h*_*p*_ and *h*_*t*_), which are projected into the joint latent space (*z*_*p*_ and *z*_*t*_). The combined loss function, ℒ_*P enCL*_, is calculated using the similarity matrices between these embeddings, incorporating both contrastive and masked language modeling losses to guide the learning process. The final transformer blocks of both pLM and bLM are fine-tuned using ℒ_*P enCL*_, alongside the projection heads *g*_*pLM*_ and *g*_*bLM*_.

The training was conducted on four NVIDIA A100 GPUs, leveraging data distributional training to facilitate a large negative batch size’s crucial factor in contrastive learning. As shown in Figure S3, the AllGather function is employed to concatenate representations from each GPU, enabling the computation of full similarity matrices across the distributed system. This approach ensures robust and scalable training, with backpropagation occurring across all GPUs using PyTorch. The latent embedding size for the protein (*z*_*p*_) and text (*z*_*t*_) joint embedding space were set to 512, with an Adam optimizer used at a base learning rate of 0.0016, adjusted according to global_batch_size*/*256 ×base_lr. The temperature parameter for both ℒ_*GC*_ and ℒ_*P F C*_ was set to 0.8, and dropout within the projection heads was set to 0.1. Sequence lengths were capped at 512 tokens for text and 1024 tokens for proteins, including special start and end tokens. The *h*_*p*_ and *h*_*t*_ embeddings were sized at 1280 and 768, respectively, with input dimensions for the projection heads adjusted accordingly. The training utilized floating-point 16 precision.

Our final PenCL model, used for benchmarking evaluations, was pretrained for 20 epochs, with the 5-epoch checkpoint employed for design experiment evaluations. The validation set was created from a 20% random split of the combined Swiss-Prot and Pfam datasets, enabling comprehensive evaluation across diverse protein families and textual descriptions.

This comprehensive pretraining strategy integrates multiple loss objectives and distributed computing techniques to effectively train PenCL, creating a biologically meaningful and semantically rich representation space that bridges protein sequences and textual descriptions.

##### A.2.3 PenCL Ablation Pretraining Analysis

To evaluate the impact of different loss components in the PenCL model, we conducted a series of ablation experiments. Each ablation modifies the pretraining loss terms to assess their individual contributions to the model’s performance. The ablations and their pretraining setups are detailed as follows:

###### No ablation

The full loss function (ℒ_*P enCL*_) is employed, combining the Global Contrastive loss (ℒ_*GC*_), the Protein Family Contrastive loss (ℒ_*P F C*_), and the masked language modeling losses for both text (ℒ_*bML*_) and protein sequences (ℒ _*pML*_). This configuration leverages the full SwissProt and Pfam datasets by sampling pairs from both databases and fine-tuning the language models in PenCL model.

###### Ablation 1

The Protein Family Contrastive loss (ℒ_*P F C*_) is removed, setting its contribution to zero while retaining the other loss terms (ℒ_*GC*_ + ℒ_*bML*_ + ℒ_*pML*_). This setup still samples pairs from both SwissProt and Pfam, thus utilizing the full dataset, and fine-tunes the language models in the PenCL model. This configuration tests the importance of homology-based learning by excluding explicit family contrastive learning.

**Figure S2:**
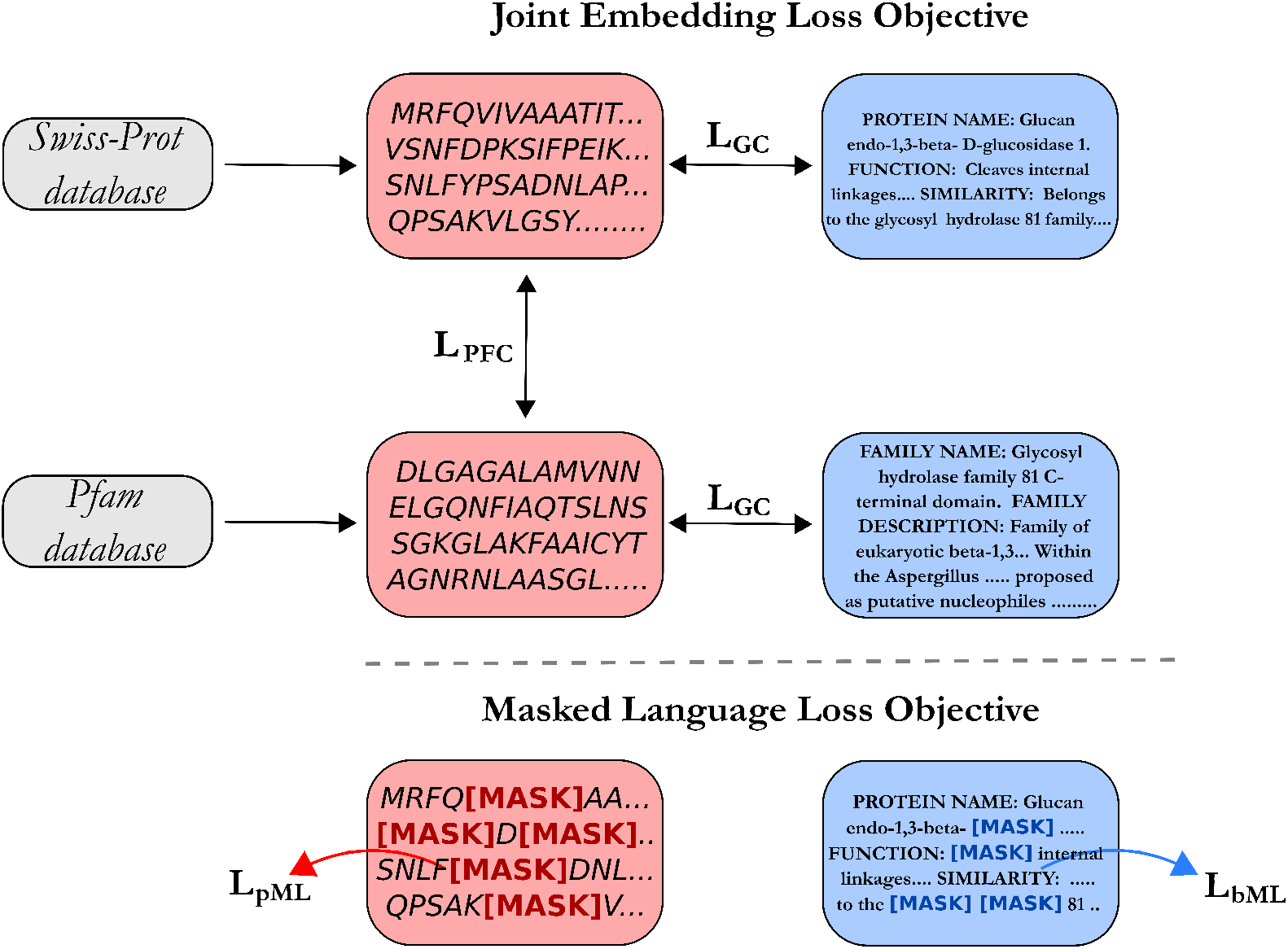
Overview of PenCL’s joint embedding loss and masked language loss objectives. The figure demonstrates the data flow from Swiss-Prot and Pfam databases into the contrastive learning and masked language modeling components of the model. Losses ℒ_*GC*_, ℒ_*P F C*_, ℒ_*bML*_, and ℒ_*pML*_ are computed to align protein sequences and their descriptions within the joint embedding space.

###### Ablation 2

The Protein Family Contrastive loss (ℒ_*P F C*_) is removed, setting its contribution to zero while retaining the other loss terms (ℒ_*GC*_ + ℒ_*bML*_ + ℒ_*pML*_). Unlike other setups, this ablation only samples from the SwissProt database and does not utilize Pfam data, limiting the homology context and relying mainly on cross-modal alignment. The mask *m* in Supplementary Algorithm 2 is not applied since Pfam database is not sampled alongside the Swissprot database. The language models in the PenCL model is still fine-tuned, including the language models.

###### Ablation 3

This configuration retains only the Global Contrastive loss (ℒ_*GC*_), excluding the Protein Family Contrastive and masked language modeling losses. It samples data from SwissProt database and fine-tunes the language models in the PenCL model, focusing on cross-modal protein-text alignment without leveraging homology or intra-modal context.

###### Ablation 4

The final ablation retains the Global Contrastive loss (ℒ_*GC*_) but with a distinct configuration indicated by the symbols in the figure, possibly modifying the contrastive alignment strategy. Unlike other ablations, this setup does not fine-tune the language models, and the remaining training is restricted to the projection heads. This setup explores the effect of alignment learning without further adjusting the pretrained language models.

These ablations are designed to probe the significance of particular loss components and the impact of sampling strategies on model performance, and expose the need for balanced cross-modal and intra-modal objectives in the absence of extensive protein-text corpora. PenCL performance under these ablations in a remote homology task is reported in Table S7.

###### Algorithm 1

Stage 1 Dataloader for PenCL Pretraining

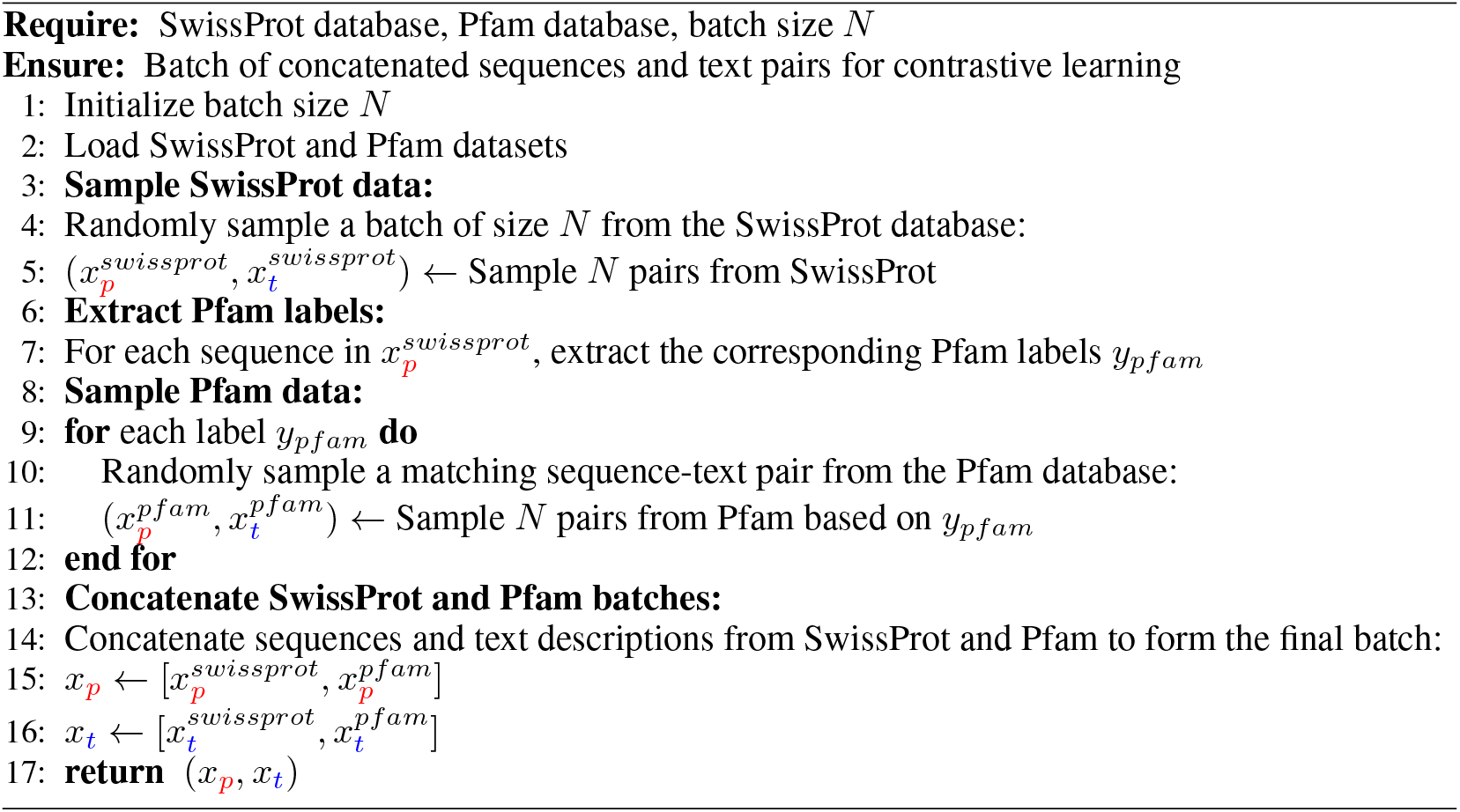

#### A.3 Stage 2: Improving text prompt conditioning with Facilitator module

##### A.3.1 Facilitator Architecture and Loss

The second stage of the PenCL model incorporates an additional module known as the Facilitator, designed to further refine the alignment of text embeddings with protein embeddings generated by PenCL. This Facilitator module, originally introduced by ProteinDT [23] and shown to empirically improve the generative stage, is a simple autoencoder that improves text embeddings by reconstructing the protein embeddings. Unlike ProteinDT, which employs mean-square error (MSE) loss, we utilize a Maximum Mean Discrepancy (MMD) loss to enhance the alignment between protein sequence representation (*z*_*p*_) and the augmented text representation (*z*_*c*_ = *f*_Facilitator_(*z*_*t*_)). This loss is defined as follows:

###### Stage 2 Pretraining Loss

Stage 2 Pretraining Loss

The Stage 2 training loss objective for the Facilitator is given by:

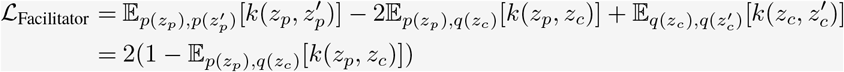

where we compute the maximum mean discrepancy (MMD) loss between the protein sequence representation *z*_*p*_ and the produced protein sequence representation (i.e., *z*_*c*_ = *f*_Facilitator_(*z*_*t*_)), *p*(*z*_*p*_) is the distribution over *z*_*p*_, *q*(*z*_*c*_) is the distribution over *z*_*c*_, and *k*(·, ·) is a Gaussian kernel,

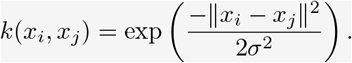

and *σ* is a hyperparameter. The simplification after the second equality results from the fact that the first and third terms evaluate to unity.

Over the training epochs, both the training and validation losses for MMD consistently decrease without evident signs of overfitting, while the MSE loss shows slight overfitting towards the later epochs (Fig. S4A). This suggests that MMD loss is more robust in aligning the text and protein embeddings without the risk of overfitting, which is particularly beneficial in a multi-modal setting.

###### Algorithm 2

PenCL’s main pretraining algorithm.

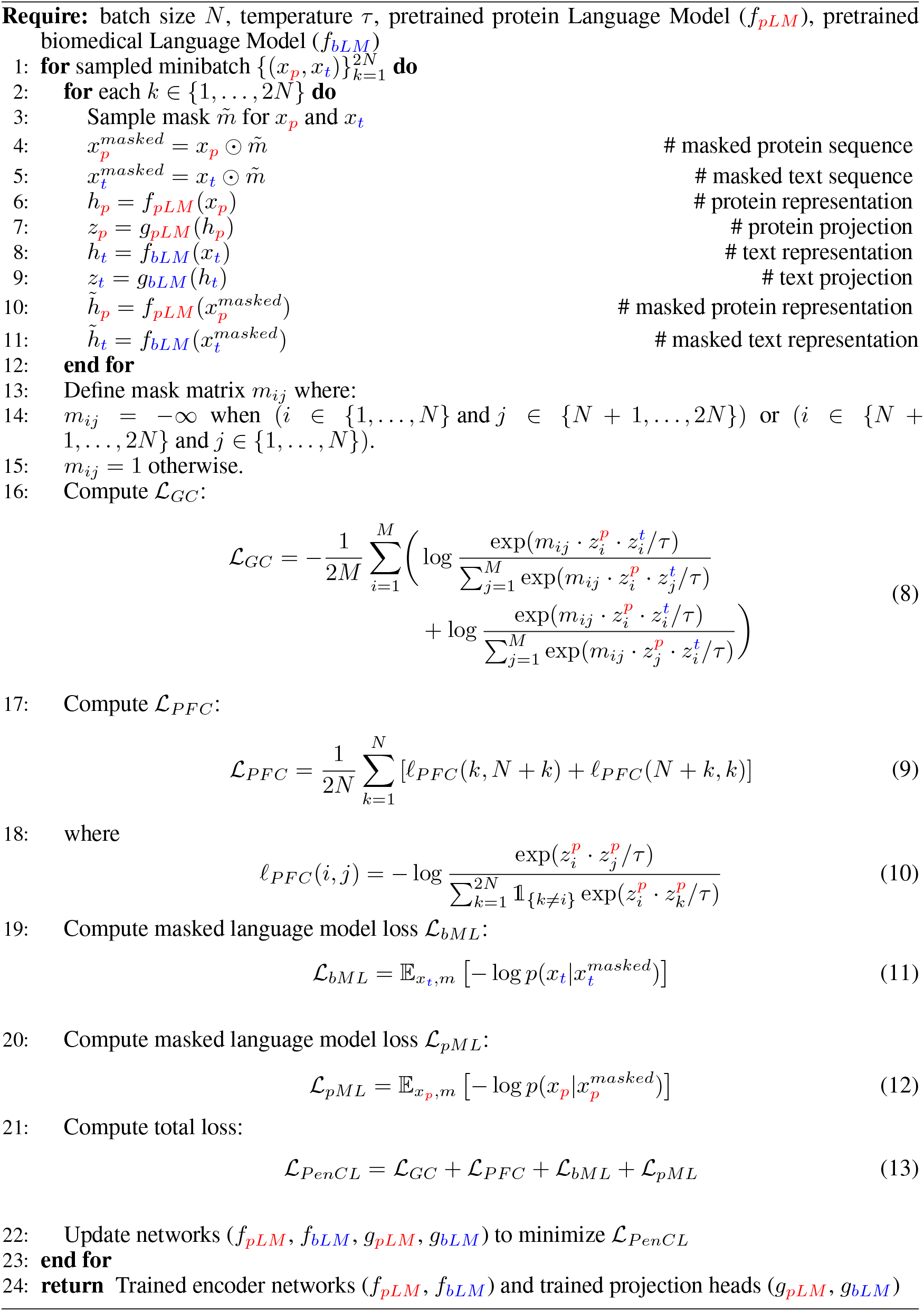

**Figure S3:**
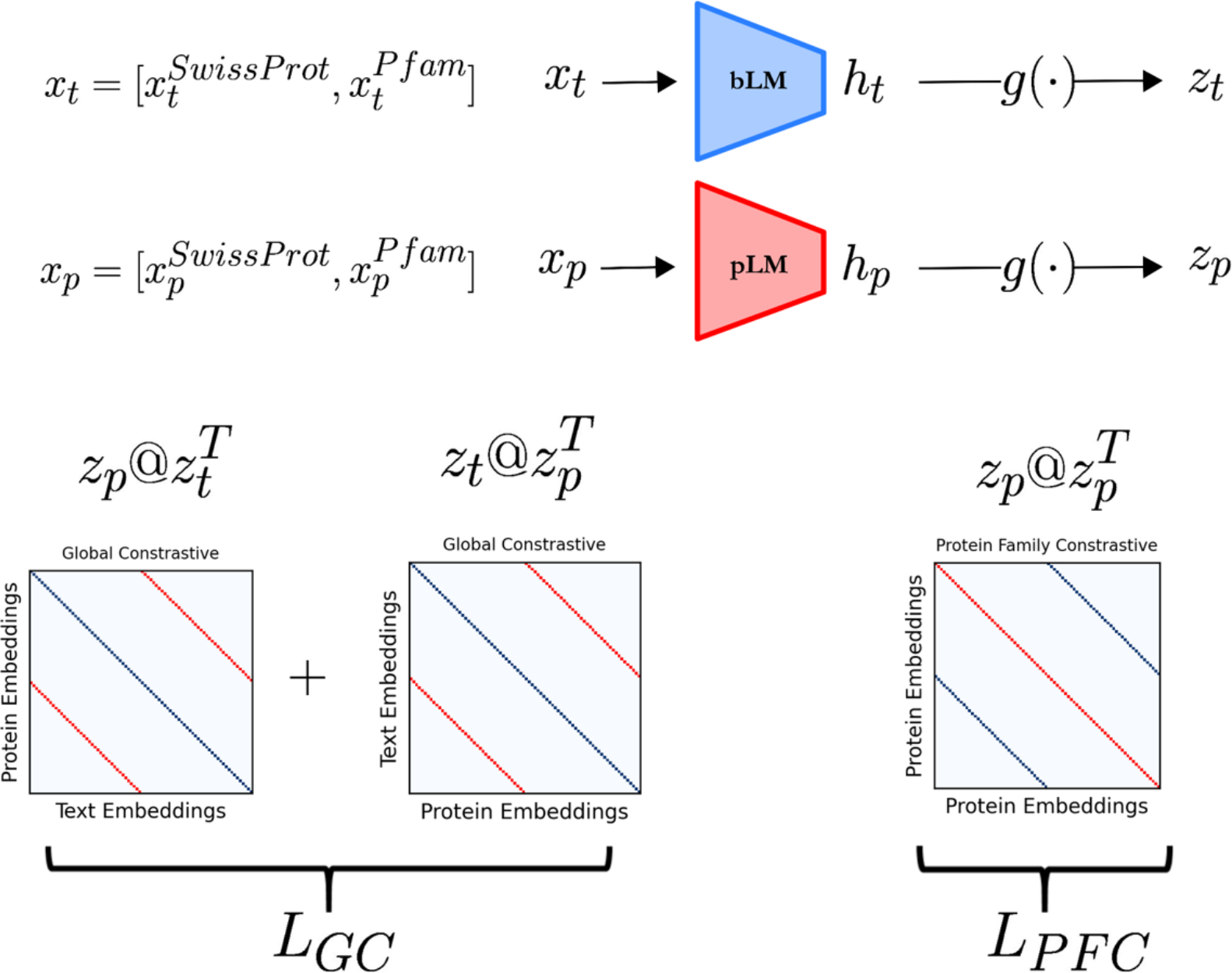
PenCL’s architecture and forward pass for similarity computation. Protein and text representations are generated by their respective language models, projected into the joint embedding space, and used to compute similarity matrices critical for defining ℒ_*GC*_ and ℒ_*P F C*_. The AllGather function is employed to concatenate representations across GPUs, ensuring robust loss computation during training.

The Facilitator employs a simple multilayer perceptron (MLP) architecture where the input and output dimensions are matched, defined as follows:

~~~
# Main neural network structure
self.main = nn.Sequential(
  weight_norm(nn.Linear(in_dim, hid_dim), dim=None), # Weight-normalized linear layer
  nn.GELU(), # GELU activation function
  nn.Dropout(dropout, inplace=True), # Dropout layer
  weight_norm(nn.Linear(hid_dim, out_dim), dim=None) # Weight-normalized output layer
)
~~~

Here, `nn.Linear` is PyTorch’s linear affine transformation, `weight_norm` applies weight normalization by decoupling the magnitude of the weight tensor from its direction, `nn.GELU()` is the non-linear activation function, and `nn.Dropout()` serves as a regularization layer to reduce overfitting. The input (`in_dim`) and output (`out_dim`) dimensions match the joint embedding space dimensions, which, in our case, is 512. The hidden dimension size (`hid_dim`) is 1024.

The training process was configured with specific hyperparameters tailored to optimize the performance of the Facilitator module. We set the validation size to 0.2, with a batch size of 512 and 64 workers for efficient data loading. The model was trained using a single GPU with a precision of 32-bit. The dataset type used was “default”, with the model type specified as “pfam” and the loss function set to Maximum Mean Discrepancy (MMD). A dropout rate of 0.0 was employed, and the learning rate was set at 1e-3, optimized using the Adam optimizer. We trained two versions of the Facilitator: one for the benchmarked PenCL model, which was pretrained for 20 epochs, and another for the finetuned SH3 experimental design model, trained for 15 epochs. This configuration ensured that each Facilitator was specifically adapted to its respective model context, enhancing the alignment between the protein and text embeddings for their intended applications.

##### A.3.2 Facilitator Pretraining

The pretraining of the Facilitator involves fine-tuning the autoencoder to improve the alignment between the text and protein embeddings. Supplementary Algorithm 3 outlines the pretraining protocol for the Facilitator module.

###### Algorithm 3

Stage 2 Pretraining of Facilitator

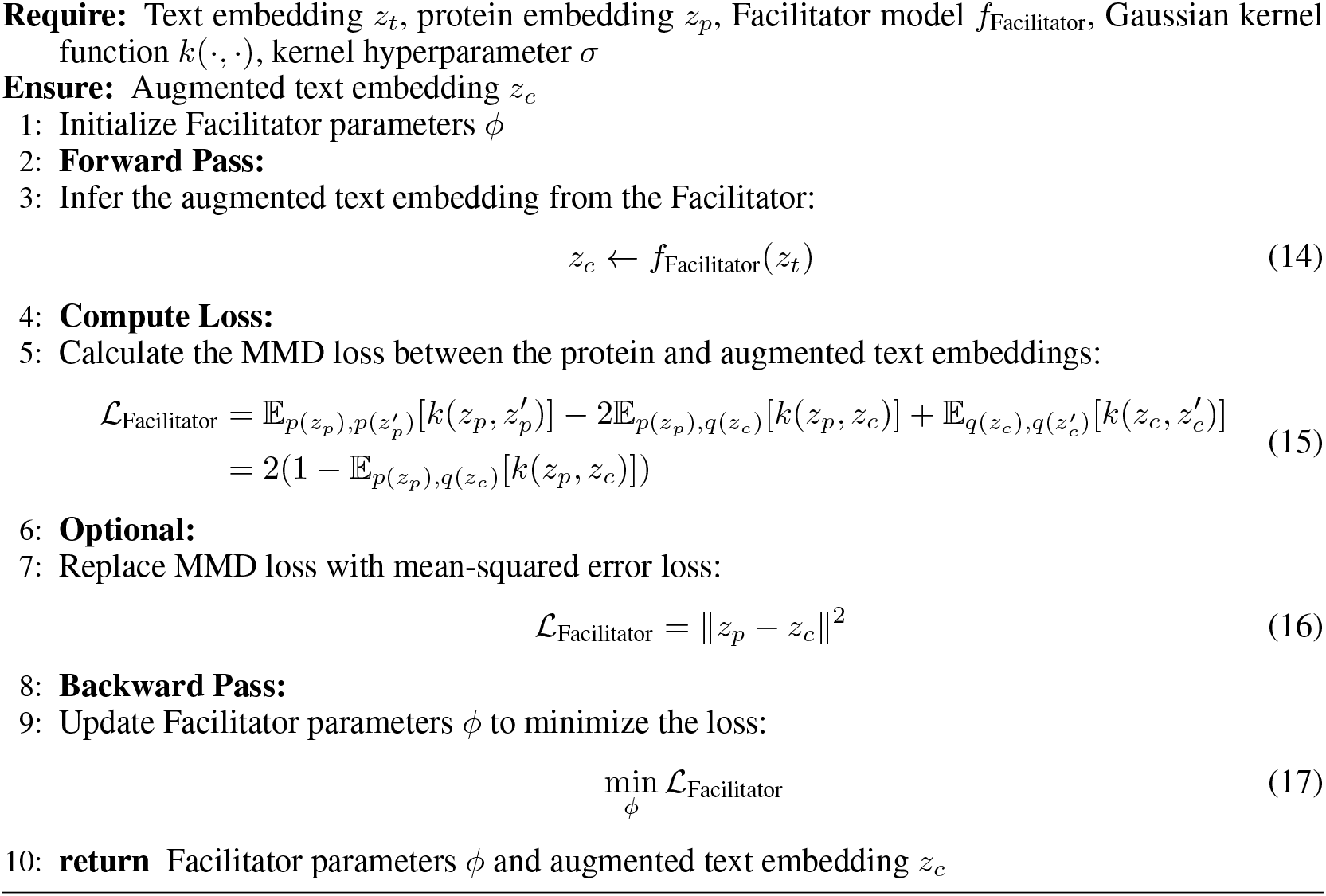

This training process ensures that the Facilitator refines the text embeddings by aligning them more closely with their corresponding protein embeddings, enhancing the overall performance of the PenCL model. Empirically, we found that improved agreement between the norm of text embedding ||*z*_*t*_|| and protein embedding ||*z*_*p*_|| induced by the facilitator was a key factor in improving overall performance of the BioM3 model (Fig. S4B).

#### A.4 Stage 3: Order-Agnostic AutoRegressive Diffusion Model with Text Description Conditioning (ProteoScribe)

##### A.4.1 Architecture and Loss of ProteoScribe

In Stage 3, we employ an order-agnostic autoregressive diffusion model (ARDM) to generate artificial protein sequences guided by textual descriptions, referred to as ProteoScribe (Figure S5). Unlike previous approaches such as ProteinDT [23], which use autoregressive or discrete diffusion decoders, our ARDM implementation allows flexible order-agnostic sequence generation or in-painting of (non-contiguous) motifs. Inspired by EvoDiff [24], which first applied ARDM to proteins, our model further incorporates text-prompted conditionals, enhancing the capacity to generate sequences based on text inputs.

The architecture of ProteoScribe integrates an efficient transformer [67, 68, 75] with several optimizations to handle long sequences efficiently. The backbone of the model is an efficient transformer with linear attention, allowing it to scale linearly with sequence length. Key components include:

**Figure S4:**
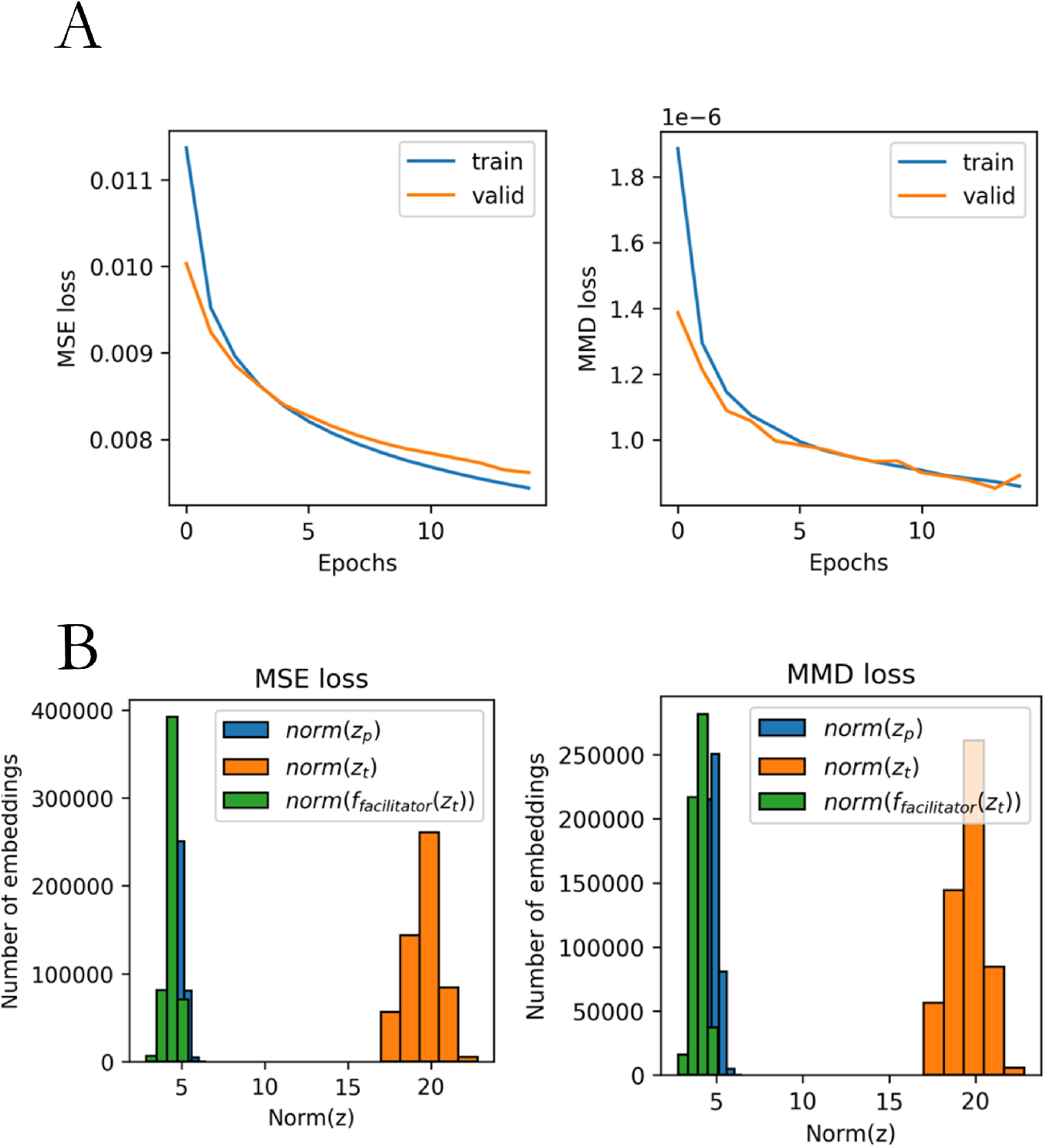
Performance of the Facilitator Module in PenCL Stage 2 Pretraining. (A) Comparison of training and validation loss curves for the MSE (left) and MMD (right) losses over epochs. The MMD loss demonstrates a more stable reduction with less overfitting after 10 epochs compared to the MSE loss, indicating better alignment performance between text and protein embeddings.(B) Distribution of embedding norms for protein embeddings (‖ *z*_*p*_ ‖), text embeddings (‖ *z*_*t*_ ‖), and augmented text embeddings produced by the Facilitator (‖*z*_*c*_‖= ‖ *f*_Facilitator_(*z*_*t*_) ‖). The Facilitator effectively matches the norms of the text and protein embeddings, enhancing the compatibility and alignment between the modalities, and improving overall performance of the BioM3 model.

- **Sinusoidal Time Embeddings**: These embeddings encode diffusion steps, rescaled to match the training step scale (total diffusion steps *D* = 1024), providing unique encoding for each diffusion time point, which is critical for tracking sequence degradation and subsequent reconstruction.
- **Axial Positional Embeddings**: A technique used in machine learning, particularly in transformer-based models, to encode positional information in a more efficient and scalable manner than traditional positional embeddings.
- **Transformer Blocks**: The transformer consists of 16 blocks, each with 16 attention heads and an embedding dimension of 512, resulting in a model with approximately 90M parameters. This configuration provides a context window of 1024, matching the diffusion length.

**Figure S5:**
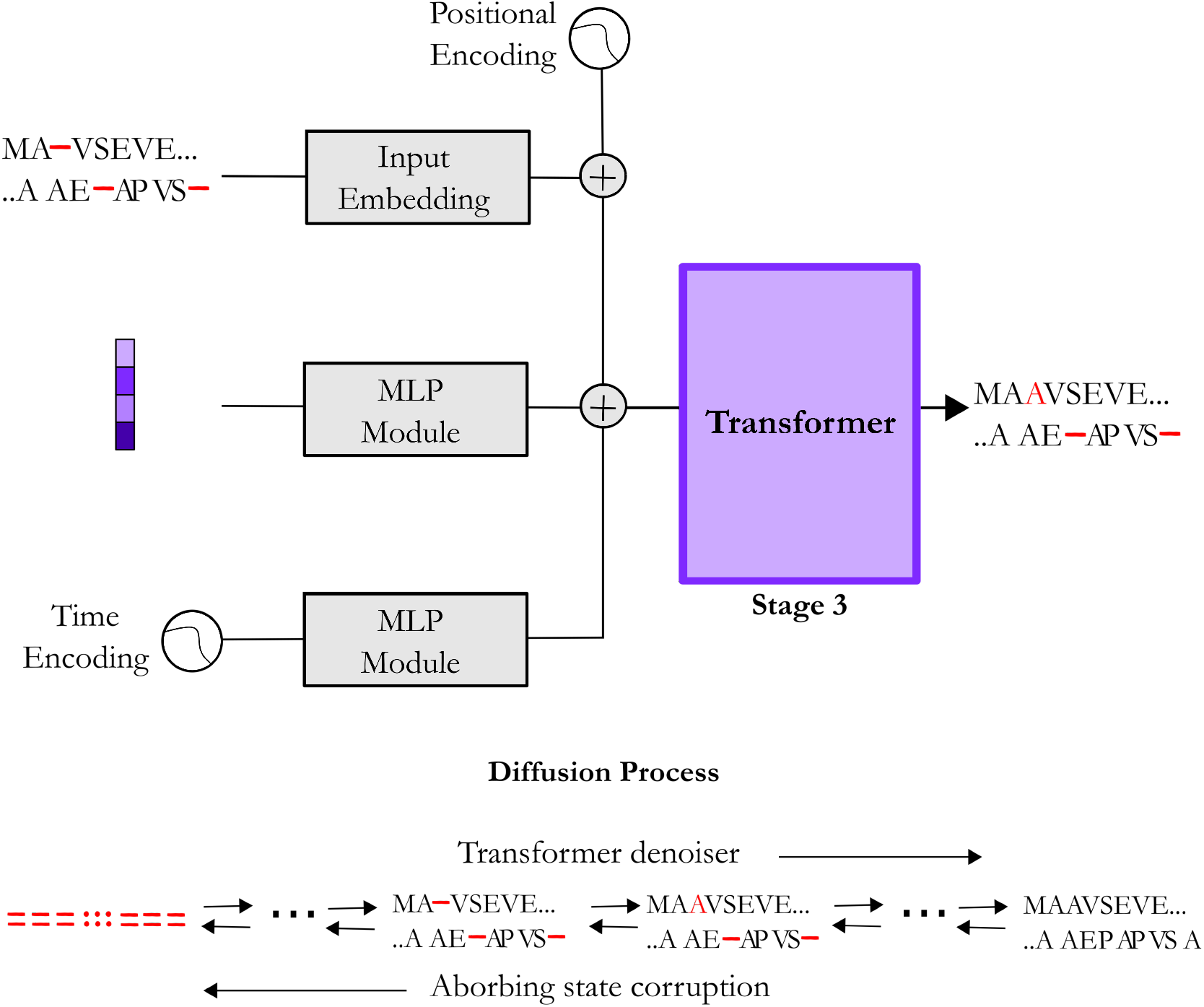
Schematic Illustration of Text-guided ProteoScribe. Illustration of the conditioned transformer-based architecture pretrained with an order-agnostic autoregressive diffusion process for proteins (ProteoScribe). This allows for conditioned embeddings inferred from text prompts and achieves order-agnostic autoregressive in-painting.

Sequences are trained to a length of 1022, with the start and end tokens appended, and padding added if sequences are shorter than 1024.

- **Input Embeddings**: We employ a token embedding layer with 29 input token representations, including 20 amino acids, 3 special tokens, 5 alphabetic special tokens found in the UniProt database, and 1 absorbing state placeholder token.
- **Multilayer Perceptron (MLP) Embeddings**: An MLP architecture with a structure of a linear layer, softmax activation, and another linear layer is used to embed the time and augmented text latent embeddings *z*_*c*_, crucial for integrating information from the text prompt into the sequence generation process.

The training loss objective for ProteoScribe is defined by an order-agnostic objective function, allowing the model to reconstruct sequences in any arbitrary order, enhancing the flexibility of sequence generation:

###### Stage 3 Pretraining Loss

The pretraining loss objective for Stage 3, referred to as ProteoScribe, employs a text-guided autoregressive diffusion model. This objective combines diffusion-based noise prediction, autoregressive sequence modeling, and contrastive learning, and is formally defined as follows:

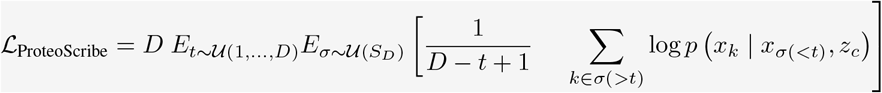

where *D* is the number of timesteps, *t* is the timestep, 𝒰 (*S*_*D*_) is the uniform distribution of all possible orderings of positions *σ* ∈*S*_*D*_, *x* is the protein sequence, *x*_*k*_ is the *k*^th^ token of the protein sequence, *x*_*σ*(*<t*)_ is the set of (possibly non-contiguous) tokens specified or decoded prior to time *t*, and *z*_*c*_ = *f*_Facilitator_(*z*_*t*_) is the protein sequence representation produced by the Facilitator.

##### A.4.2 ProteoScribe Pretraining

The pretraining of ProteoScribe involves fine-tuning the discrete diffusion model to generate sequences conditioned on text prompts. The training protocol consists of sampling time points, corrupting sequences at these points using absorbing states and reconstructing the sequence by conditioning on the remaining uncorrupted sequence and the text-derived embedding *z*_*c*_ as detailed in Supplementary Algorithm 4. This approach enables the model to flexibly adapt the reconstruction order, allowing in-painting of specific sequence motifs or regions based on the conditioning context and leverages the order-agnostic nature of the ARDM while expanding generation capabilities by enhancing latent variable conditioning of augmented text embedding *z*_*c*_. This allows for flexible sequence reconstruction driven by the text prompt. This capability makes ProteoScribe a powerful tool for generating sequences that align closely with natural language descriptions, opening new avenues in protein design and annotation tasks.

###### Algorithm 4

Stage 3 Pretraining Protocol for ProteoScribe

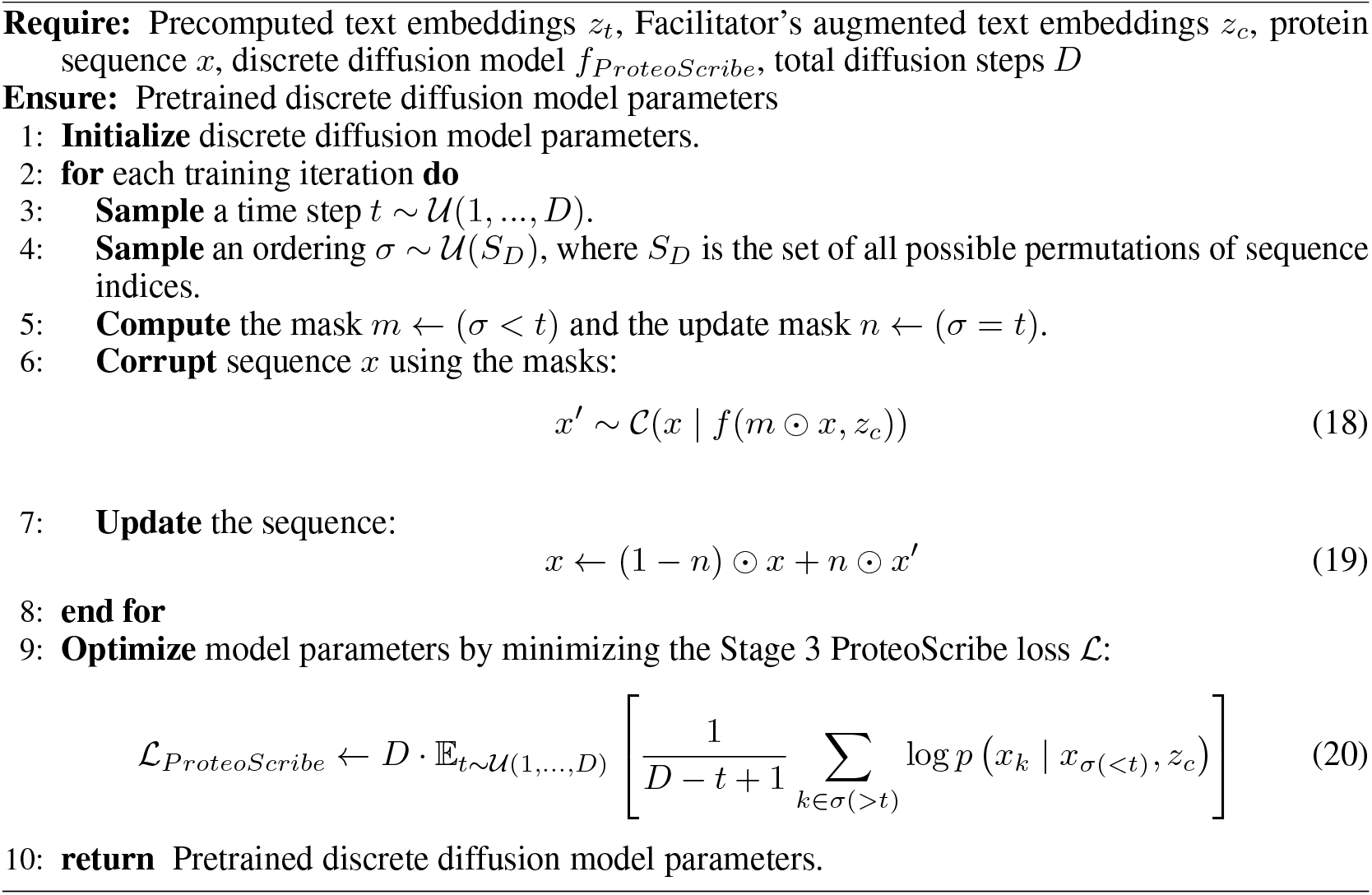

The experimental design model of ProteoScribe was pretrained using an AdamW optimizer with a learning rate of 1 ×10^−4^, weight decay of 1× 10^−6^, and a warmup period of 500 steps. This training was conducted over 100 epochs on an 80/20 random split of the SwissProt dataset, utilizing a local batch size of 32 distributed across 4 NVIDIA A100 GPUs with the ZeRO optimizer (DeepSpeed Stage 2), ensuring efficient scaling and resource utilization. This model configuration was primarily used for experimental design applications.

Additionally, a second model, with identical architecture and parameter settings, underwent a more extensive pretraining regimen. Initially, it was trained for 600 epochs on the SwissProt split dataset. Following this, the optimal checkpoint – selected based on performance on the validation set – was further fine-tuned on a combined 80/20 split of the SwissProt and Pfam datasets. This additional phase consisted of 3 million training steps, equivalent to approximately 10 epochs. This two-phase pretraining approach was designed to enhance the model’s performance for *in silico* benchmarking and evaluation tasks, leveraging the expanded data set to capture a broader representation of protein sequences.

##### A.4.3 Sampling and In-painting with ProteoScribe

After pretraining ProteoScribe, the model can be used to generate new protein sequences guided by text prompts or to in-paint missing motifs in existing sequences. The generation process leverages the augmented text embeddings *z*_*c*_, derived from the Facilitator, to condition the ProteoScribe ARDM during the sampling process.

###### Sampling New Sequences

The process of sampling new sequences given a text prompt is detailed in Supplementary Algorithm 5. Here, the generation starts by initializing the sequence *x* with a start token. The ordered denoising positions, denoted by *σ*, can either be predefined, randomly sampled, or uniformly sampled from all possible permutations of sequence indices. During each time step, the model generates the next token by conditioning on the already generated parts of the sequence and the Facilitator embedding *z*_*c*_. This iterative denoising process continues until the entire sequence is completed, resulting in a sequence that aligns with the provided text prompt.

###### In-painting Motifs

ProteoScribe can also be utilized for in-painting motifs within sequences, as shown in Supplementary Algorithm 6. In this scenario, a specific motif is provided at predefined positions within the sequence. The starting time step *t*_start_ controls when the in-painting process begins, and the ordered denoising positions *σ* determine where the model will perform decoding. The process ensures that the known motif remains fixed, while the surrounding sequence is generated to fill in any missing regions, creating a complete, coherent sequence. This flexibility to specify motif positions and adjust the denoising trajectory makes ProteoScribe particularly powerful for targeted sequence design.

Both algorithms highlight the versatility of ProteoScribe in generating novel protein sequences and filling in gaps with biologically relevant motifs. The ability to control denoising positions either randomly or through user-defined settings offers significant flexibility for various protein design applications, providing precise control over how sequences are generated or modified.

#### A.5 Finetuning Model Details and Experimental Protocol for SH3 Design

##### A.5.1 Details and Preparation of the SH3 Dataset

To fine-tune the ProteoScribe model for the design and experimental testing of SH3 domains, we curated a comprehensive SH3 dataset consisting of text-protein pairs from three primary sources, each matched with the Pfam label PF00018. The first source, SwissProt [38], provided 599 protein-text pairs, representing multimeric proteins that include SH3 domains within their original, natural context. These sequences typically reflect the full-length proteins where SH3 domains are naturally embedded, capturing the broader functional landscape of SH3 domains as integral parts of larger protein complexes. The second source, the Pfam database [38], contributed 16,566 protein-text pairs focusing specifically on isolated SH3 domains rather than full-length proteins. This dataset offers a broad representation of SH3 domain sequences, including variations and homologs across different organisms, but lacks the surrounding protein context found in multimeric forms. Additionally, we included 7,865 protein-text pairs consisting of isolated SH3 domains [15]. For this dataset, text prompts were constructed by retrieving the paralog name (`SH3 PARALOG NAME`) and the functional annotation description (`PARALOG FUNCTION`) associated with each domain. An example of a constructed prompt is:

###### Algorithm 5

Sampling New Sequences with ProteoScribe

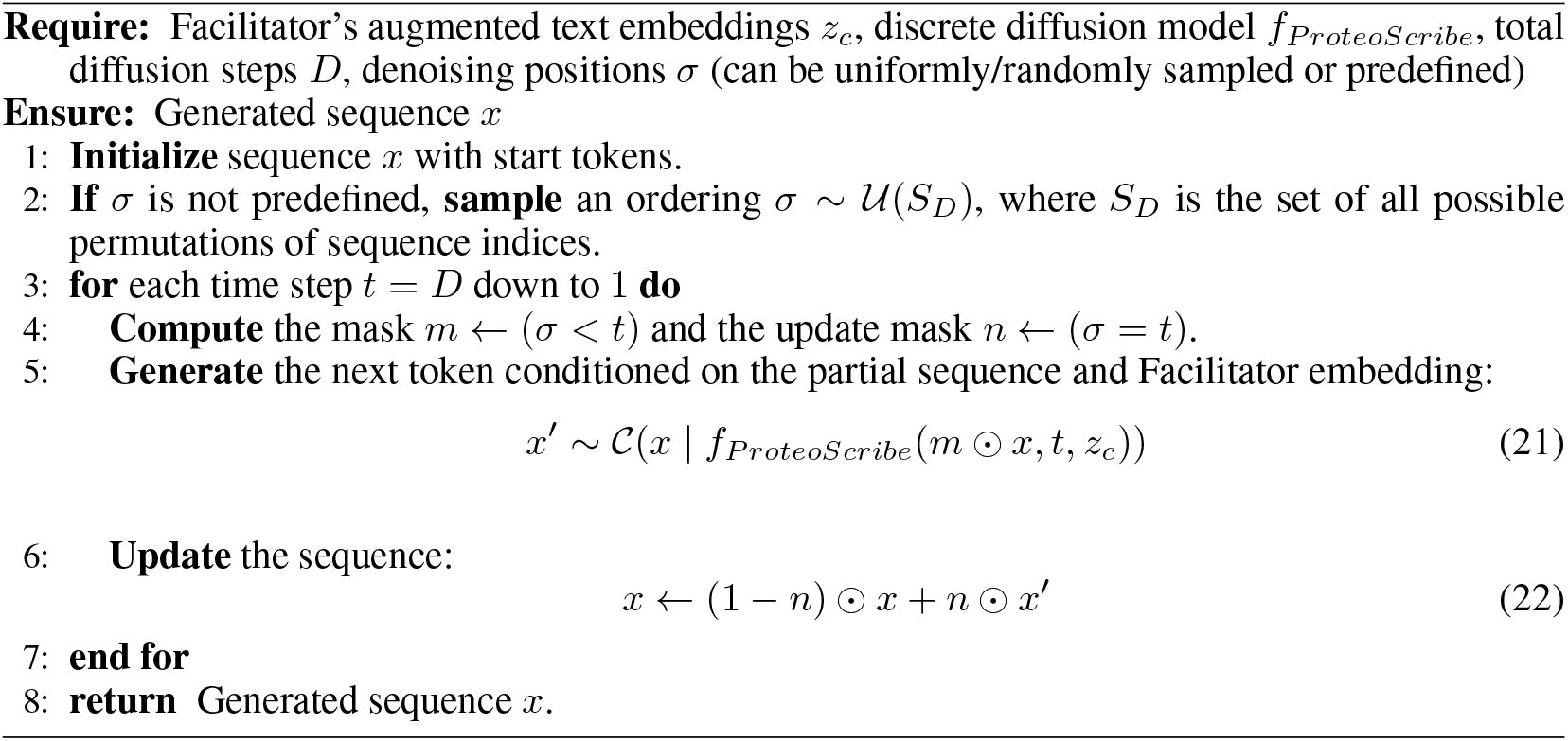

###### Algorithm 6

In-painting Motifs or Scaffolds in a Sequence with ProteoScribe

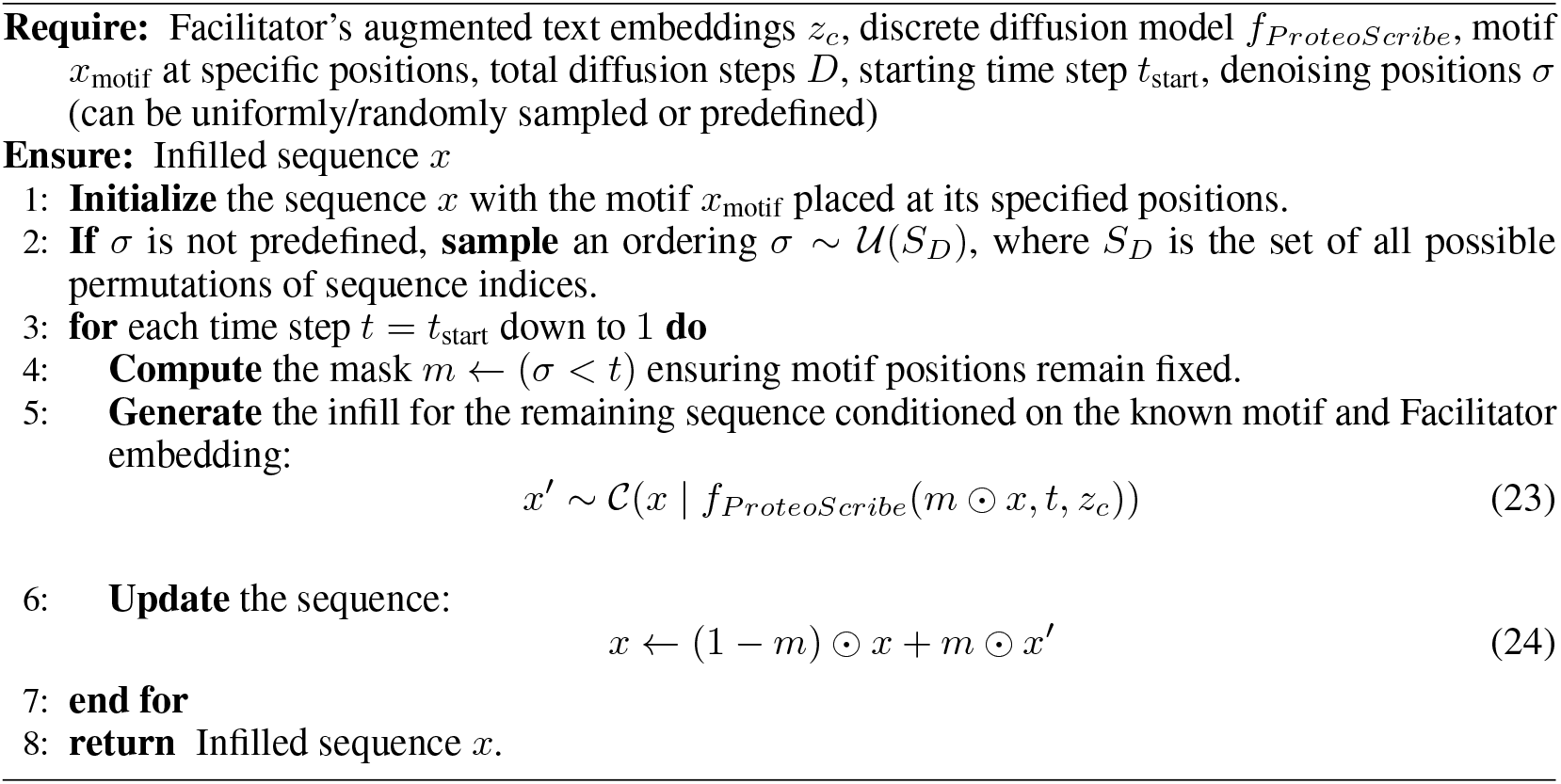

> PROTEIN NAME: SH3 domain. LINEAGE: The organism lineage is cellular organisms; Eukaryota; Opisthokonta; Fungi; Dikarya; Ascomycota; Saccharomyceta; Saccharomycotina; Saccharomycetes; Saccharomycetales; Saccharomycetaceae; Torulaspora; Torulaspora delbrueckii. SH3 PARALOG NAME: BEM1. PARALOG FUNCTION: Protein containing SH3-domains; involved in establishing cell polarity and morphogenesis; functions as a scaffold protein for complexes that include Cdc24p, Ste5p, Ste20p, and Rsr1p.

The combined dataset was split into an 80/20 training-validation ratio to ensure robust model finetuning and evaluation. Sequences longer than 1,022 amino acids were excluded to conform to the sequence length constraints of the ProteoScribe model. This curated SH3 dataset provides a diverse set of sequences and functional descriptions, allowing the model to learn and generalize the intricate relationships between SH3 domains and their corresponding textual annotations. By including multimeric proteins from SwissProt, which reflect the natural contextual embedding of SH3 domains, along with isolated domains from the Pfam database and other sources, the dataset enhances the model’s ability to capture both the specific functional features of SH3 motifs and their roles within larger protein assemblies.

##### A.5.2 SH3 Finetuning of ProteoScribe

The finetuning of ProteoScribe for SH3 domain design involved utilizing the pretrained Stage 1 PenCL model (epoch 5 checkpoint) and the corresponding Stage 2 Facilitator checkpoint. Both the PenCL and Facilitator models were kept frozen during finetuning, maintaining their pretrained weights. The embeddings *z*_*t*_ and *z*_*c*_ were inferred directly from the text prompts associated with the SH3 domains, allowing the model to leverage the learned representations from the initial stages without further modification.

The finetuning process followed the same training hyperparameters and architectural settings as described in Appendix A.4.1, ensuring consistency across stages. The training was conducted with a local batch size of 32 over 1,000 epochs using 4 NVIDIA A100 GPUs. However, it is important to note that the hardware does not necessarily need to be A100 GPUs specifically; any CUDA-compatible GPUs with sufficient VRAM and support for deep learning software and tools can be utilized for this process. Distributed training was implemented using the DeepSpeed Stage 2 ZeRO optimizer, facilitating efficient multi-GPU scaling and memory management.

The final model used for SH3 design was selected based on the checkpoint that demonstrated optimal validation loss without overfitting, which occurred at epoch 53. This strategy ensured that the model retained the ability to generalize to new SH3 sequences while leveraging the rich contextual information encoded during the pretraining phases. The fine-tuned model thus integrates the text-protein alignment capabilities of the PenCL and Facilitator stages with the specialized knowledge of SH3 domain structures and functions, providing a robust platform for experimental design.

##### A.5.3 Procedure of Sampling SH3 Designs with Five Prompts

To evaluate the effectiveness of the ProteoScribe model in generating functional SH3 domain sequences, we sampled a total of 5,000 sequences across five distinct textual prompts. These prompts were designed to explore different ways of guiding the generation of sequences that emulate the functional characteristics of Sho1^SH3^ domains. The first corresponds to the text annotation for Sho1^SH3^ that was seen during finetuning. The second and third are ablated versions of the first: the second prompt retains functional information, whereas the third includes only the name of the Sho1^SH3^ domain. The fourth and fifth prompts were entirely new: the fourth includes only the protein name, and the fifth incorporates the name and functional description.

###### SHO1 Prompt 1

**PROTEIN NAME:** SH3 domain. **LINEAGE:** The organism lineage is cellular organisms; Eukaryota; Opisthokonta; Fungi; Dikarya; Ascomycota; saccharomyceta; Saccharomycotina; Saccharomycetes; Saccharomycetales; Saccharomycetaceae; Saccharomyces; Saccharomyces cerevisiae; Saccharomyces cerevisiae S288C. **SH3 PARALOG NAME:** SHO1. **PARALOG FUNCTION:** Transmembrane osmosensor for filamentous growth and HOG pathways; involved in activation of the Cdc42p- and MAP kinase-dependent filamentous growth pathway and the high-osmolarity glycerol (HOG) response pathway; phosphorylated by Hog1p; interacts with Pbs2p, Msb2p, Hkr1p, and Ste11p.

###### SHO1 Prompt 2

**PROTEIN NAME:** SH3 domain. **SH3 PARALOG NAME:** SHO1. **PARALOG FUNC-**

**TION:** Transmembrane osmosensor for filamentous growth and HOG pathways; involved in activation of the Cdc42p- and MAP kinase-dependent filamentous growth pathway and the high-osmolarity glycerol (HOG) response pathway; phosphorylated by Hog1p; interacts with Pbs2p, Msb2p, Hkr1p, and Ste11p.

###### SHO1 Prompt 3

**PROTEIN NAME:** Src Homolog 3 (SH3) domain in the high osmolarity signaling protein SHO1. **FAMILY NAMES:** Family names are Src Homolog 3 (SH3) domain.

###### SHO1 Prompt 4

Src homology 3 (SH3) domain-containing members of the SHO1 family.

###### SHO1 Prompt 5

Key transmembrane SH3 domain protein in osmotic sensing for filamentous growth and HOG pathways, involving Cdc42p/MAP kinase interactions and phosphorylation.

A total of 1,000 sequences were generated under each prompt and structurally predicted using ColabFold with MMSeqs2 [72] for fast MSA retrieval to enhance alignment speed and maintain alignment quality. For each design, five structural predictions were generated, and the TM-score computed against the wild-type Sho1^SH3^ domain from *Saccharomyces cerevisiae*. Sequences with an average TM-score above 0.5 were considered structurally similar to the wild-type SH3 domain.

From each set of 1,000 sequences per prompt, 200 designs were randomly selected based on structural similarity and functional relevance. Specifically, sequences were required to have top BLAST hits in the NCBI non-redundant database with terms relevant to Sho1 function, including “osmosensor”, “transmembrane”, “SHO1”, “SH3”, “osmolarity”, “kinase”, and “Src Homolog 3”. Due to poor structural predictions for sequences generated from Prompt 3, which were likely caused by high novelty and diversity, the 200 designs with the highest TM-scores were selected without strict BLAST hit criteria. This resulted in fewer than 200 sequences from Prompt 3 meeting the BLAST criteria, and thus, the remaining designs to fill the quota were incorporated into the pool of Prompt 5 designs.

The final distribution of SH3 designs included 200 sequences each from prompts 1, 2, and 4; 67 sequences from prompt 3; and 333 sequences from prompt 5. Additional controls included the wild-type Sho1^SH3^ sequence from *S. cerevisiae* and a null allele, along with a calibration curve consisting of a mutation screen of the wild-type allele with various phenotypic effects [15]. In total, the final gene designs synthesized included 200, 200, 67, 200, and 317 sequences for Prompts 1, 2, 3, 4, and 5, respectively, for a total of 984 sequences across all five prompts.

While this approach specifically assesses our model’s ability to generate functional SH3 domains, the broader framework exemplifies the potential of starting from textual descriptions of desired functions to infer latent variables that guide protein generation. This demonstrates how natural language inputs can direct protein design, ultimately enabling the synthesis of proteins with enhanced functionalities based on textual and conceptual inputs.

##### A.5.4 Experimental Details of the *In Vivo* Assay

The *in vivo* assay for evaluating SH3 domain designs was based on previously published protocols [15], with specific adjustments made to suit the current study’s objectives.

###### Gene Construction

Synthetic SH3 domains were generated by reverse translating the protein sequences into DNA sequences, followed by codon optimization for yeast expression, focusing on balanced GC content and optimized codon usage for *S. cerevisiae*. The oligonucleotides corresponding to each gene were synthesized on microarray chips (Twist Inc.) and included primer annealing sites and padding sequences to ensure uniform amplification conditions. PCR amplification was conducted using KAPA HiFi polymerase with specific forward and reverse primers. The amplified products were then digested with EcoRI and BamHI, ligated into the PRS316 plasmid backbone containing a Sho1 N-terminal membrane domain, and subsequently transformed into Agilent Electrocompetent XL1-Blue cells. The cells were grown in LB media containing ampicillin, and plasmids were purified and pooled.

###### Yeast Transformation

The constructed plasmids were transformed into the haploid yeast strain SS101, built on the W303 background, featuring genetic knockouts of Ssk2 and Ssk22 to remove the Sho1-independent osmosensory pathway components. Transformation was carried out using a LiAc-PEG high-efficiency transformation protocol. Transformed yeast cells were grown in Sc-Ura media and repeatedly passaged until the transformed allele pool achieved normal growth and doubling times.

###### SH3 Domain Selection Assay

The SH3 domain selection assay followed the protocol detailed in Ref. [15], with modifications to extend the selection period to 72 hours to provide a longer observation window for assessing the functionality of each SH3 domain design *in vivo*. Cultures were grown in Sc-Ura media initially, then transferred to YPD media supplemented with KCl to apply selective pressure. This extended selection period allowed for a more thorough evaluation of the designed SH3 sequences under osmotic stress conditions. After the 72-hour selection, plasmids were extracted, and amplicon libraries were prepared for Illumina sequencing. The relative enrichment (*r*.*e*.) of each designed SH3 sequence *x* was calculated using the following expression:

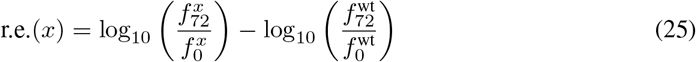

where 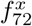 and 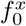 are the frequencies of observing the sequence *x* at the 72-hour and 0-hour time points, respectively, and 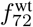 and 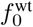 are the corresponding frequencies for the wild-type SH3 domain. This equation quantifies the enrichment of each design relative to the wild-type, providing insight into the evolutionary advantage conferred by each synthetic SH3 domain under selective conditions.

This adapted assay allowed for a comprehensive evaluation of synthetic SH3 domains over an extended selection period, offering valuable insights into the adaptability and performance of the designs in the yeast cellular environment.

##### A.5.5 Experimental Details of the *In Vitro* Assay

The *in vitro* binding assay was conducted to evaluate the binding affinity of the designed SH3 domains to their target ligand, pbs2 MAPKK, following previously established protocols [15]. The protocol involves measuring the intrinsic tryptophan fluorescence of the SH3 domain upon titration with a synthetic peptide ligand, which serves as an indicator of binding events.

###### Peptide Synthesis and Protein Expression

The pbs2 MAPKK peptides were synthesized using standard 9-fluorenylmethoxycarbonyl (Fmoc) chemistry by the Protein Chemistry Technology Center at UT Southwestern Medical Center. The peptide’s molecular mass was verified via mass spectrometry, and concentrations were confirmed by quantitative amino acid analysis.

Individual SH3 domains were cloned into the pET28b-P expression vector to generate N-terminal His6-tagged proteins. For protein expression, the plasmids were transformed into *E. coli* BL21 (DE3) cells and cultured in TB media with 50 *µ*g/mL kanamycin. Protein expression was induced with 200 *µ*M IPTG when cultures reached an OD_600_ of 0.8-1.2, followed by incubation at 18^°^C overnight. Cells were harvested, lysed, and proteins were purified using Ni-NTA affinity chromatography, followed by further purification using size exclusion chromatography. Purified SH3 proteins were aliquoted, flash-frozen using liquid nitrogen, and stored at (−80)^°^C.

###### Binding Assay Protocol

The binding affinity between Sho1^SH3^ domains and synthetic pbs2 MAPKK ligands was measured by observing the changes in intrinsic tryptophan fluorescence upon titration of the peptide ligand into a fixed concentration (0.25 *µ*M) of Sho1^SH3^ protein in HEPES buffer (20 mM HEPES, 50 mM NaCl, pH 7.3-7.6). The fluorescence measurements were performed using a Fluorolog-3 spectrofluorometer with excitation and emission wavelengths set to 296 nm and 330 nm, respectively.

###### Data Analysis

The resulting fluorescence readings were fitted to a binding curve using the equation:

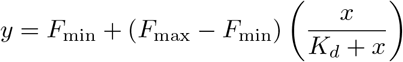

where *y* represents the fluorescence reading, *x* is the ligand concentration, *K*_*d*_ is the dissociation constant, and *F*_min_ and *F*_max_ are the minimum and maximum fluorescence values, respectively. The fitting was performed using the `scipy.optimize.curve_fit` module in Python. This approach allowed us to quantitatively assess the binding affinity of the designed SH3 domains.

### B Supplementary Results

#### B.1 Stage 1: Detailed Results and Analysis of PenCL Inter-Clan and Intra-Clan Clustering Analysis

Figure S6 presents the PenCL architecture and its performance in integrating protein sequences and text descriptions within a unified joint embedding space. The architecture utilizes two pretrained language models: the biomedical Language Model (bLM) for processing text and the protein Language Model (pLM) for processing protein sequences. Embeddings from both models are projected into a common space, facilitating cross-modal alignment through the Global Contrastive Loss (ℒ_*GC*_), which aligns correct protein-text pairs while separating unrelated pairs. The Protein Family Contrastive Loss (ℒ_*P F C*_) further refines this alignment by ensuring that sequences from the same protein family are more tightly clustered, distinguishing them from other families (Fig. S6A).

The clustering performance was assessed using the Calinski-Harabasz Index (CHI) and the Davies-Bouldin Index (DBI) [70], standard metrics for clustering quality evaluation. PenCL demonstrated significant improvements over single-modality models, including ProtT5 [29], and ESM2 [29], as indicated by higher CHI scores and lower DBI values (Fig. S6B). This evaluation was particularly relevant in inter-clan clustering, which measures the ability to separate all superfamilies within the BioM3 curated Pfam database. PenCL also showed substantial improvements in intra-clan clustering, which assesses the ability to separate protein families within a given superfamily for all superfamilies presented in the BioM3 Pfam database (Fig. S6C). Notably, PenCL outperformed the state-of-the-art multimodal language model ProtST [22], suggesting that PenCL’s enhancements stem from more than merely including the natural language modality. This underscores PenCL’s effectiveness in capturing the structural and functional nuances of protein families based on textual information.

##### Visualization of Intra-Clan and Inter-Clan Clustering of Protein-Text Samples

PenCL’s joint embedding space was visualized using PCA projections to evaluate the model’s capacity to distinguish between inter-clan (superfamily) and intra-clan (family) clusters (Fig. S7). The interclan clusters represent broader superfamily classifications, while intra-clan clusters correspond to specific protein family groupings. The separation of these clusters within the PCA space demonstrates PenCL’s ability to capture hierarchical relationships within protein families, informed by both sequence and text data.

Figure S7 shows distinct PCA clusters for various protein families, such as PF01926 and PF00009, illustrating how PenCL effectively maps homologous sequences and their corresponding text prompts into similar regions of the embedding space. This alignment not only facilitates the clustering of proteins within the same family but also enhances the interpretability of the model’s embeddings in representing the functional and structural diversity of protein sequences.

##### Benchmarking Performance in Zero-Shot and Remote Homology Detection Tasks

PenCL’s performance was further evaluated in zero-shot learning tasks for subcellular localization and enzyme classification using text-guided protein retrieval (Table S5). PenCL’s multimodal embeddings allowed it to perform competitively with ProtST [22] in predicting subcellular localization and enzyme function without task-specific fine-tuning, using prompts previously defined for ProtST. This suggests that further performance improvements could be achieved through prompt engineering specifically tailored for PenCL. The best-performing text annotation recovered by PenCL achieved an accuracy of 0.402 for subcellular localization and 0.531 for enzyme classification, highlighting the impact of well-crafted text prompts.

In remote homology detection tasks (Table S6), PenCL significantly outperformed traditional sequence comparison methods like BLASTp [71] and multimodal models such as ProtST [22]. Homology detection accuracy was assessed at multiple levels: fold, superfamily, and family. PenCL achieved the highest top 1 and top 5 accuracy scores across these levels, particularly excelling in family-level retrieval with top 1 accuracy of 0.977 and top 5 accuracy of 0.987. These results underscore PenCL’s superior ability to detect distant homologs, which is critical in protein engineering and functional annotation. PenCL also outperformed state-of-the-art single-modality language models such as ProtT5 [29] and ESM2 [26], further highlighting the robust representations learned by PenCL.

Ablation studies conducted on the remote homology detection task revealed that ℒ_*P F C*_ played a crucial role in enhancing performance (Table S7), particularly in distinguishing closely related families. This finding emphasizes the importance of aligning text and protein embeddings at both a global and family-specific level, demonstrating how PenCL’s design facilitates nuanced understanding of homology and functional relationships across diverse protein sequences.

**Figure S6:**
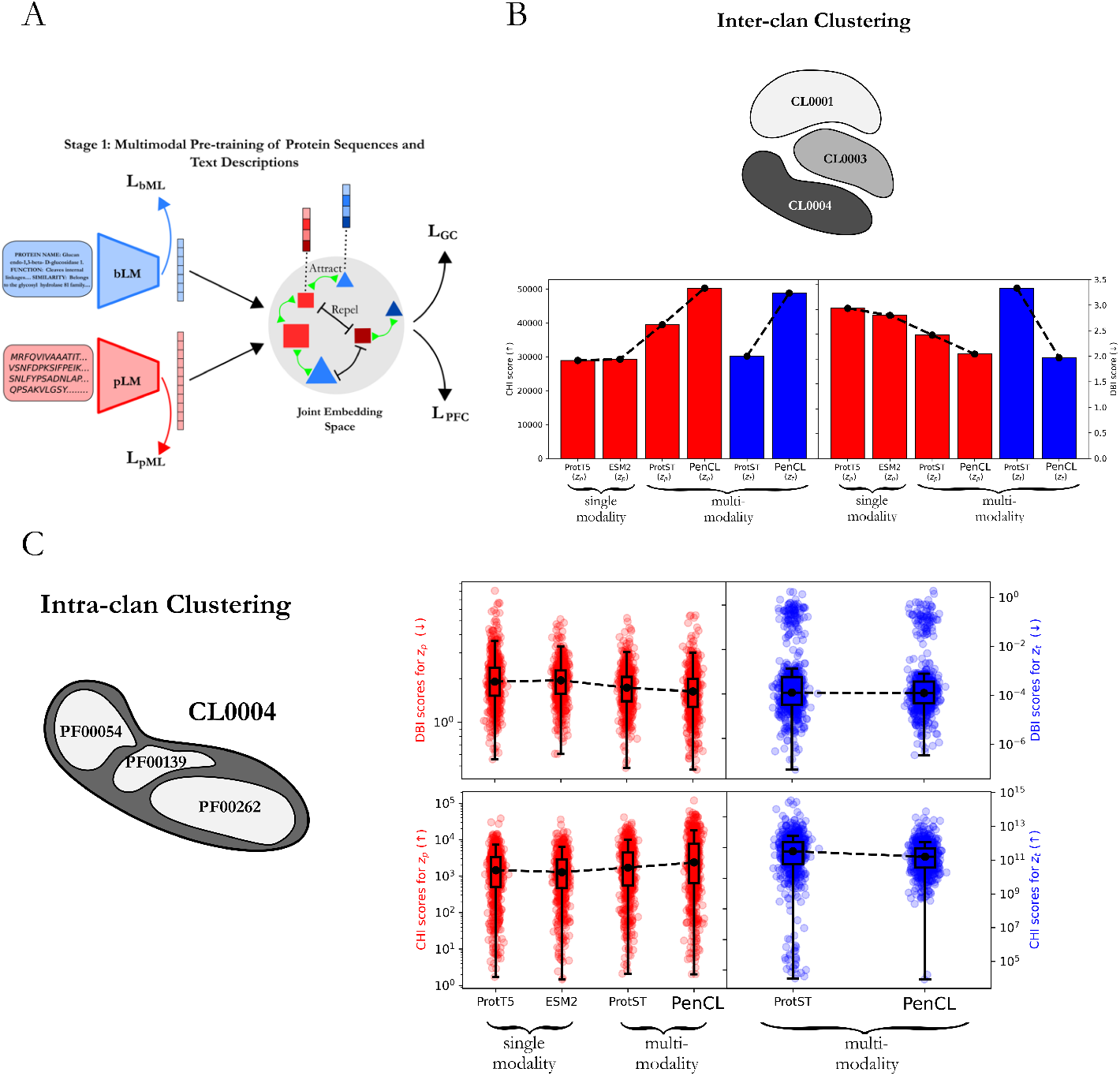
Multimodal Pretraining of Protein Sequences and Text Descriptions in Stage 1. (A) Illustration of the PenCL architecture integrating protein sequences and text descriptions. The biomedical Language Model (bLM, blue) processes text, and the protein Language Model (pLM, red) processes sequences. The joint embedding space combines these, with triangles representing text and squares representing sequences. Different colors and sizes indicate different protein families and textual descriptions. Global Contrastive loss (ℒ_*GC*_) aligns correct text-sequence pairs and separates incorrect ones, while Protein Family Contrastive loss (ℒ_*P F C*_) aligns homolog pairs from the same family and separates those from different families. More details of the losses in Stage 1 are in the Appendix A.2. (B-C) Comparison of clustering performance of PenCL against single modality models (ProtT5 [29], ESM2 [26]) and multimodal models (ProtST [22]) for both inter-clan (superfamily) and intra-clan (protein family) clustering. Performance is measured using the Calinski-Harabasz Index (CHI) and Davies-Bouldin Index (DBI). Higher CHI and lower DBI values indicate better performance. Red bars show clustering based on protein embeddings, and blue bars show clustering based on text embeddings. Overall, the intra-clan and inter-clan clustering metrics show that multimodal models like ProtST and PenCL elevate the interpretation of protein sequences and improve the disentangling of clans and protein families over state-of-the-art single-modality models like ProtT5 and ESM2, with PenCL showing improvement over ProtST.

##### Interpretation and Implications

The detailed results of PenCL Stage 1 pretraining and benchmarking highlight the advantages of integrating multimodal protein-text representations. By aligning sequence and textual information, PenCL not only improves clustering quality and functional annotation but also enhances the interpretability of relationships between protein families. These findings underscore PenCL’s potential to expand the toolkit of computational protein engineering, offering a robust approach for understanding and manipulating protein functions through the use of natural language text prompts.

**Table S5:** Multimodal benchmarking performance using zero-shot text retrieval for subcellular localization and enzyme classification tasks. For each task, a protein sequence is supplied and matched against a set of text prompts describing various subcellular locations or enzyme reactions, with the highest dot product score determining the predicted class. This task was defined by ProtST [22]. The table shows accuracy scores for predicting subcellular localization and classifying enzymes. **Green** indicates the top performer and **yellow** indicates the second-best performer for each task.

| Model | Subcellular Localization | Enzyme Classification |
| --- | --- | --- |
| **PenCL (ours)** best prompt | 0.402 | 0.531 |
| **PenCL (ours)** average score across prompts | 0.2705 | 0.506 |
| ProtST | 0.42 | 0.3 |

**Table S6:** Performance comparison of different models on a zero-shot homology retrieval task. The table shows accuracy scores for retrieving sequences with homologous folds, superfamilies, and families. Accuracy is measured by whether the correct sequence with the homologous structure was retrieved evaluated across different classification levels (fold, superfamily, family) and for the top 1 and top 5 predictions. **Green** indicates the top performer, while **yellow** corresponds to the second-best performer for each metric.

| Model | Fold |  | Superfamily |  | Family |  |
| --- | --- | --- | --- | --- | --- | --- |
|  | Top 1 | Top 5 | Top 1 | Top 5 | Top 1 | Top 5 |
| BLASTp | 0.0292 | 0.0613 | 0.175 | 0.272 | 0.942 | 0.954 |
| ProtT5 (cosine similarity) | 0.085 | 0.160 | 0.168 | 0.282 | 0.659 | 0.759 |
| ProtT5 (Euclidean distance) | 0.136 | 0.228 | 0.391 | 0.512 | 0.949 | 0.972 |
| ESM2 (cosine similarity) | 0.00975 | 0.0487 | 0.00319 | 0.0175 | 0.0173 | 0.0763 |
| ESM2 (Euclidean distance) | 0.130 | 0.227 | 0.382 | 0.513 | 0.933 | 0.9615 |
| ProtST | 0.0794 | 0.160 | 0.376 | 0.511 | 0.903 | 0.958 |
| **PenCL (ours)** | 0.149 | 0.248 | 0.586 | 0.721 | 0.977 | 0.987 |

**Table S7:** Performance comparison of ablated PenCL variants on the zero-shot homology retrieval task. Details of the four ablations are provided in Appendix A.2.3. The table shows accuracy scores for retrieving sequences with homologous folds, superfamilies, and families. Accuracy is measured for top 1 and top 5 predictions across different classification levels. **Green** indicates the top performer, while **yellow** corresponds to the second-best performer for each metric. Ablation analyses were conducted with model checkpoints at 5 epochs, in contrast to the 20 epochs used in benchmarking evaluations, and, as such, the numerical values differ slightly for the “No Ablation” case from those reported in Table S6.

| Model | Fold |  | Superfamily |  | Family |  |
| --- | --- | --- | --- | --- | --- | --- |
|  | Top 1 | Top 5 | Top 1 | Top 5 | Top 1 | Top 5 |
| No Ablation | 0.149 | 0.252 | 0.589 | 0.723 | 0.973 | 0.990 |
| Ablation 1 | 0.0724 | 0.153 | 0.342 | 0.522 | 0.922 | 0.971 |
| Ablation 2 | 0.0905 | 0.180 | 0.364 | 0.525 | 0.939 | 0.976 |
| Ablation 3 | 0.0752 | 0.149 | 0.314 | 0.486 | 0.917 | 0.966 |
| Ablation 4 | 0.105 | 0.171 | 0.356 | 0.477 | 0.891 | 0.935 |

**Figure S7:**
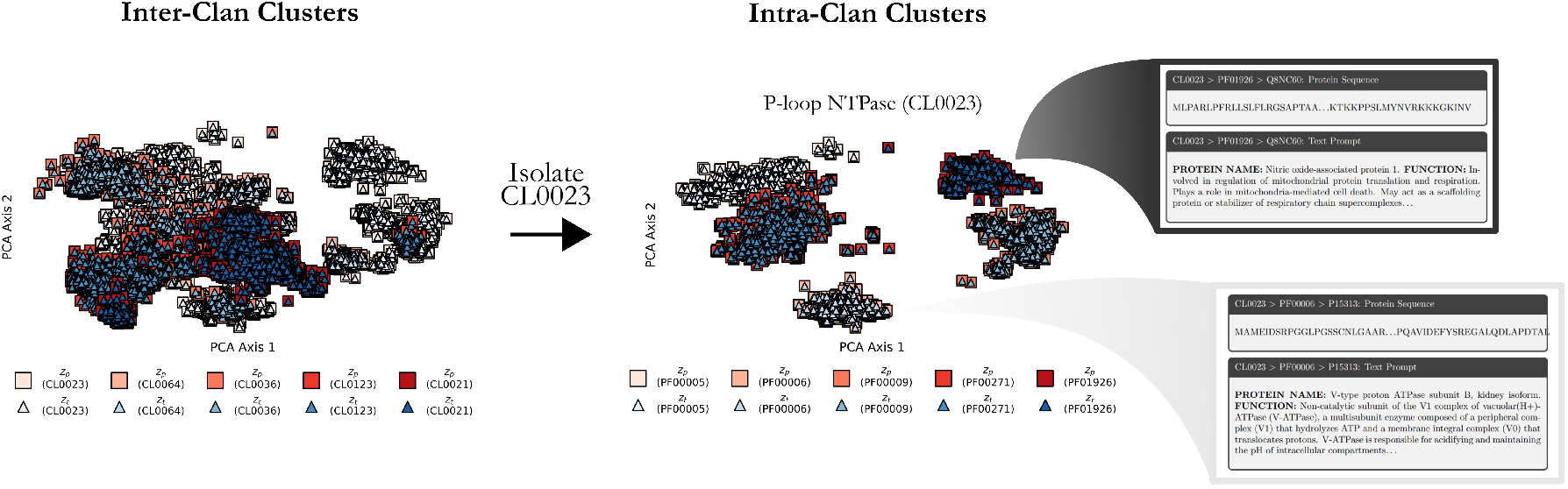
Visualization of the Hierarchical Multimodal Representations for Protein Families and Text Descriptions. PCA visualization of protein-text embeddings. The five most abundant superfamilies are shown in 2D space (Inter-Clan Clusters), and the most abundant clan (CL0023) is isolated for protein family clustering (Intra-Clan Clusters). Examples of sequences and textual descriptions are provided for families PF01926 and PF00009. The PCA visualization shows the alignment between bLM and pLM within the joint embedding space, using text-sequence pairs from training/validation splits. Text embeddings *z*_*t*_ are represented by triangles and sequence embeddings *z*_*p*_ by squares. Various clans and families are distinguished by color. These visualizations demonstrate the enhanced alignment of protein-text embeddings achieved through the PenCL model, highlighting its efficacy in capturing the relationships within and between protein families.

#### B.2 Stage 2: Results and Capabilities

##### B.2.1 Facilitator Module Performance and Analysis

Figure S4 illustrates the training and validation performance of the Facilitator module, which plays a key role in aligning text and protein embeddings in Stage 2 of PenCL. The Facilitator employs a Maximum Mean Discrepancy (MMD) loss to refine text embeddings *z*_*t*_ by augmenting them into *z*_*c*_ = *f*_Facilitator_(*z*_*t*_), aligning them closely with the protein embeddings *z*_*p*_. This alignment is crucial for enhancing text conditioning in the subsequent generative stage.

###### Training and Validation Loss Comparison

In Figure S4A, we compare the MSE and MMD losses over training epochs. The MMD loss consistently shows a slightly better alignment performance, with a smooth reduction in both training and validation losses and minimal overfitting beyond 10 epochs compared to the MSE loss. This suggests that MMD loss is more robust for maintaining the alignment quality between text and protein embeddings without compromising the generalization ability of the Facilitator.

###### Norm Matching of Embeddings

One of the primary functions of the Facilitator is to better match the norms of the embeddings between *z*_*p*_ and *z*_*t*_. As shown in Figure S4B, the distribution of embedding norms demonstrates that the Facilitator effectively aligns the norm of the augmented text embeddings ||*z*_*c*_|| = ||*f*_Facilitator_(*z*_*t*_)|| closer to that of the protein embeddings ||*z*_*p*_||. This norm matching indicates a more coherent integration of text and protein modalities, enhancing the compatibility of the augmented embeddings with the generative model’s requirements.

###### Empirical Justification for Facilitator Integration

The incorporation of the Facilitator module, introduced in ProteinDT [23], was motivated by empirical observations that addition of this module led to significant improvements in downstream protein design tasks. By improving the alignment and norm matching of embeddings, the Facilitator strengthens the overall text conditioning process, making the augmented embeddings more effective when used in the PenCL generative decoder.

These results affirm the Facilitator’s critical role in Stage 2, as it not only enhances the alignment between text and protein embeddings but also ensures that the embeddings maintain the necessary properties for high-quality generative performance in PenCL.

#### B.3 Stage 3: Results and Capabilities

In this section, we provide an in-depth analysis of the Stage 3 results from ProteoScribe, where the model is used to generate novel protein sequences conditioned on text prompts. The effectiveness of ProteoScribe in generating structurally plausible proteins is demonstrated through a combination of sequence generation, motif conditioning, and scaffold in-painting strategies, all guided by distinct textual inputs.

Figure S5 illustrates the model architecture of ProteoScribe, a transformer-based order-agnostic autoregressive diffusion model. The architecture leverages both time and positional encodings alongside the input sequence embeddings, allowing for flexible and controlled sequence in-painting. The figure highlights the key stages of the diffusion process, where sequences are incrementally corrupted and reconstructed, capturing the model’s ability to flexibly generate or repair sequence motifs based on the provided text descriptions.

##### Text-Guided Sequence Generation with Structural Validation

Figure S8 aims to demonstrate the model’s ability to generate plausible protein sequences validated by in silico predictions from five text prompts drawn from the SwissProt training data as an in-distribution test of its capacity to recapitulate protein sequence and structure. The figure showcases ProteoScribe’s capability to generate a diverse set of protein sequences with varying lengths, ranging from 72 to 617 amino acids. The designed proteins exhibit a high degree of novelty, with sequence identities to the nearest natural proteins spanning from 53.59% to 81.43%. The structural predictions, generated using ColabFold with MMSeqs2 [72] for sequence alignment, are compared against experimentally solved or predicted structures from the AlphaFold2 database. Structural alignment metrics, such as TMscore and RMSD, indicate strong structural agreement with corresponding natural proteins, demonstrating the model’s ability to maintain structural integrity while exploring novel sequence space.

##### Text-Guided Sequence Generation for Motif-Scaffold Problem

Figure S9 highlights the motif conditioning and scaffold in-painting capabilities of ProteoScribe. Using the compact calmodulin protein (PDB:1PRW) as a baseline, the figure demonstrates how the model preserves the functional motif (depicted in grey) while generating diverse scaffold designs conditioned on textual prompts. The three scaffold designs (Replicas 1, 2, and 3) maintain close RMSD values for the functional motif, confirming the model’s precise control over motif preservation. The generated scaffolds exhibit significant structural divergence, as evidenced by the RMSD values ranging from 2.587-5.437 Å, and sequence identities to 1PRW from 34.21-45.45%. These results underscore the model’s capability to generate structurally sound and novel protein designs that align closely with the guided textual inputs. The text prompt used to guide the scaffold generation was as follows:

> PROTEIN NAME: Calmodulin. FUNCTION: Calmodulin acts as part of a calcium signal transduction pathway by mediating the control of a large number of enzymes, ion channels, aquaporins and other proteins through calcium-binding. Calcium-binding is required for the activation of calmodulin. Among the enzymes to be stimulated by the calmodulin-calcium complex are a number of protein kinases, such as myosin light-chain kinases and calmodulin-dependent protein kinase type II (CaMK2), and phosphatases. Together with CCP110 and centrin, is involved in a genetic pathway that regulates the centrosome cycle and progression through cytokinesis. Is a regulator of voltage-dependent L-type calcium channels. Mediates calcium-dependent inactivation of CACNA1C. Positively regulates calcium-activated potassium channel activity of KCNN2. Forms a potassium channel complex with KCNQ1 and regulates electrophysiological activity of the channel via calcium-binding. Acts as a sensor to modulate the endoplasmic reticulum contacts with other organelles mediated by VMP1:ATP2A2. SUBUNIT: Interacts with CEP97, CCP110, TTN/titin and SRY. Interacts with MYO5A and RRAD (By similarity). Interacts with USP6; the interaction is calcium dependent (By similarity). Interacts with CDK5RAP2. Interacts with SCN5A. Interacts with RYR1 and RYR2 (By similarity). Interacts with FCHO1. Interacts with MIP in a 1:2 stoichiometry; the interaction with the cytoplasmic domains from two MIP subunits promotes MIP water channel closure. Interacts with ORAI1; this may play a role in the regulation of ORAI1-mediated calcium transport. Interacts with SYT7 (By similarity). Interacts with MYO10 and MYO1C. Interacts with SLC9A1 in a calcium-dependent manner (By similarity). Interacts with HINT1; interaction increases in the presence of calcium ions (By similarity). Interacts with HINT3 (By similarity). SUBCELLULAR LOCATION: Cytoplasm. PTM: Phosphorylation results in a decreased activity. MISCELLANEOUS: This protein has four functional calcium-binding sites. SIMILARITY: Belongs to the calmodulin family. LINEAGE: The organism lineage is Eukaryota, Metazoa, Chordata, Craniata, Vertebrata, Euteleostomi, Mammalia, Eutheria, Laurasiatheria, Artiodactyla, Ruminantia, Pecora, Bovidae, Bovinae, Bos. FAMILY NAMES: Family names are EF-hand domain pair.

Overall, these results demonstrate ProteoScribe as a robust tool for text-guided protein design, allowing for the generation of sequences with high structural plausibility and significant functional relevance. The alignment between predicted and natural structures, coupled with the ability to maintain specific functional motifs, emphasizes the utility of ProteoScribe in protein engineering and synthetic biology applications. Further experimental validation of these designed sequences, particularly focusing on their functional activities in biological systems, could provide additional insights into the practical applicability of this approach.

**Figure S8:**
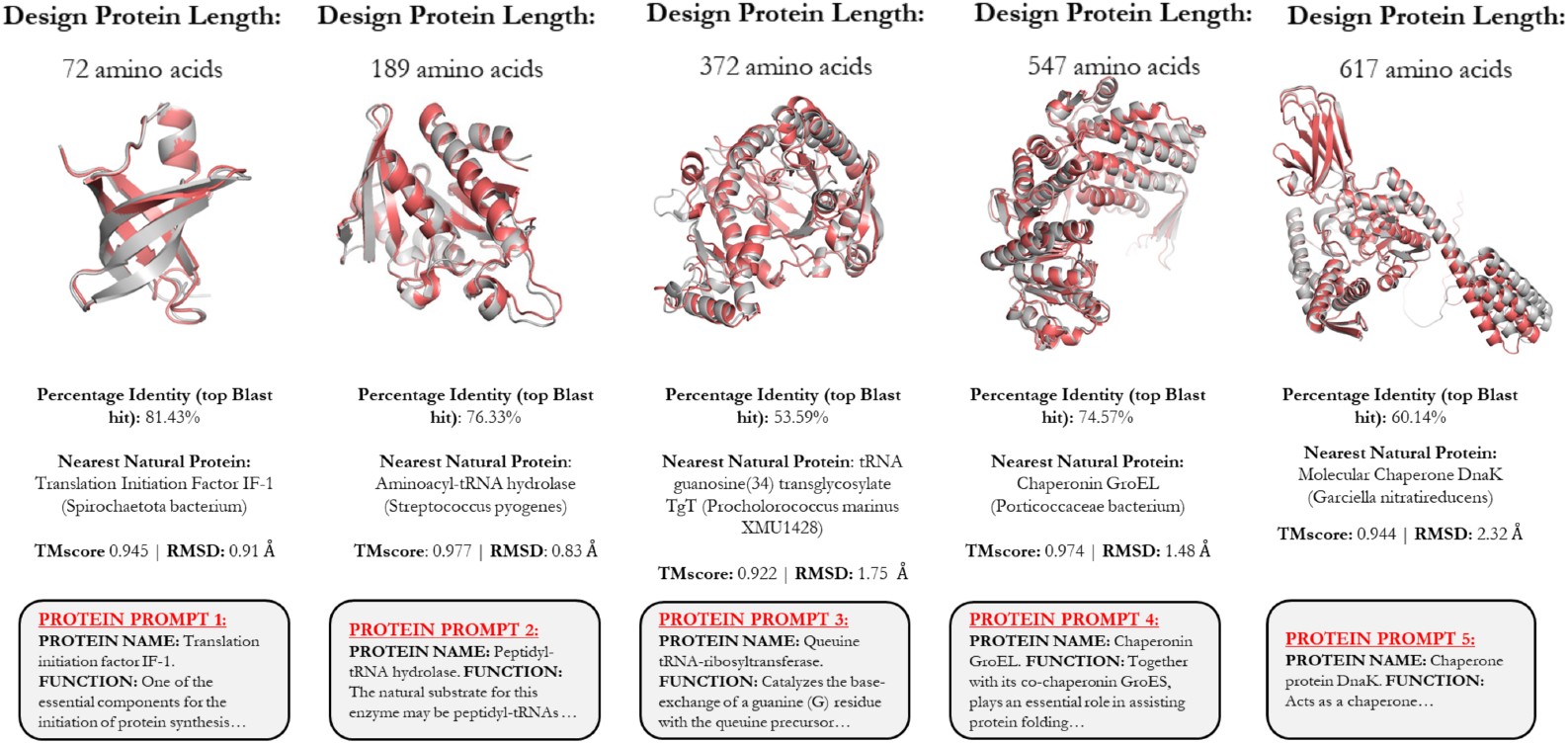
Pretrained Text-guided ProteoScribe Generates Novel and Structurally Plausible Proteins. Demonstration of ProteoScribe’s ability to generate plausible sequences based on in-distribution text annotations of proteins drawn from the SwissProt training data for sequences of various lengths (72 to 617 amino acids). The text prompt corresponding to each generated sequence is shown in the grey boxes. The nearest sequence was retrieved using BLASTp, with the percentage identity and corresponding protein name displayed. The structure of the artificial sequence (red) is predicted with ColabFold [72], and compared with the experimentally solved or predicted structure retrieved from the AlphaFold2 database [73] of the natural protein (grey) corresponding to the text prompt. Structural agreement between the natural and designed sequences is computed and displayed as TMscore and root-mean-square-distance (RMSD).

#### B.4 Experimental Validation of Prompt Engineering

To further elucidate the performance and characteristics of our text-guided protein design approach for the *Sho*1^*SH*3^ domain, we conducted additional analyses and experiments. These supplementary results provide deeper insights into the reproducibility, fitness distributions, sequence novelty, biochemical properties, and evolutionary context of our designed proteins. We present three supplementary figures and two supplementary tables that complement and extend the findings discussed in the main text.

**Figure S9:**
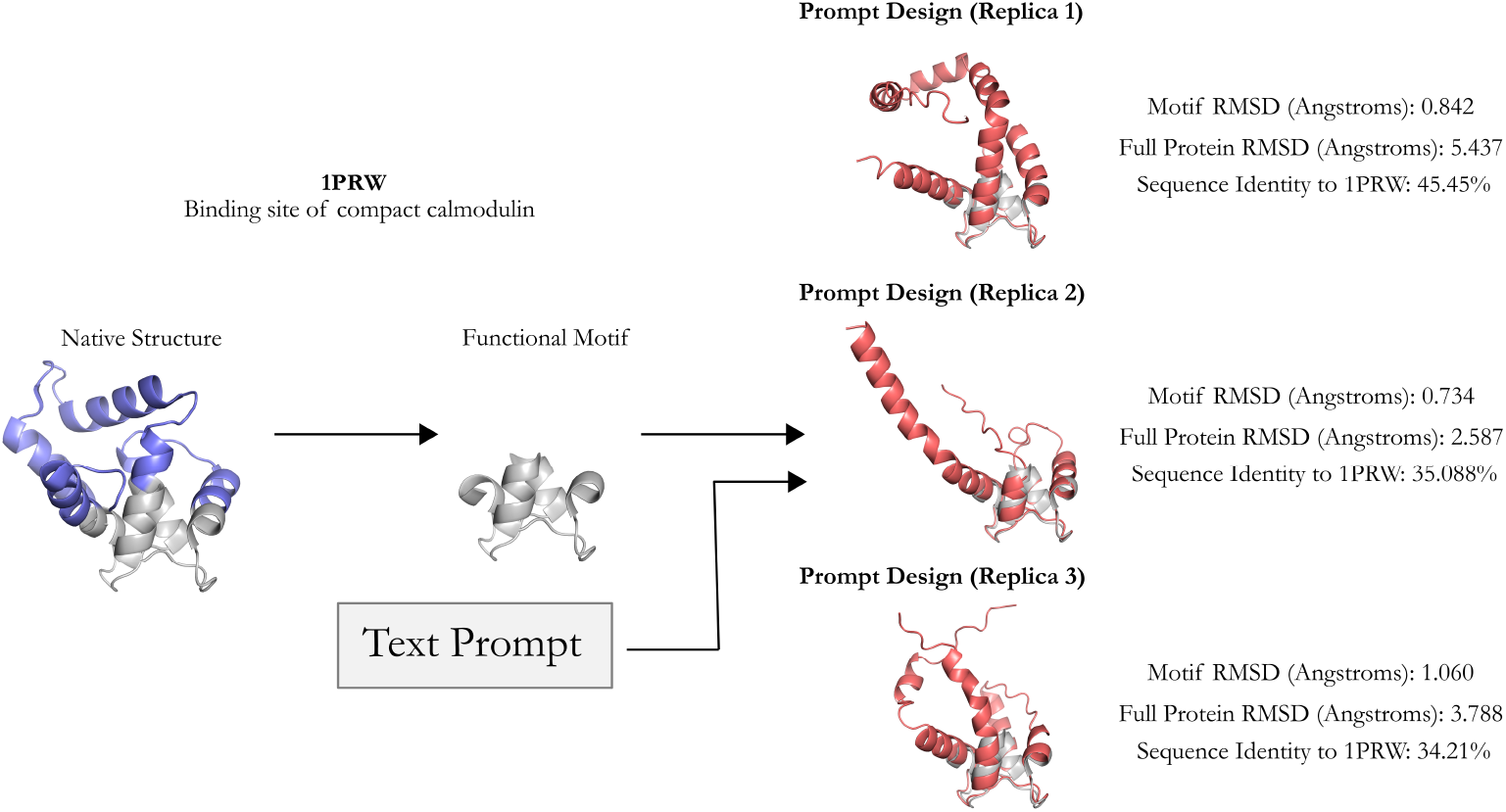
llustration of Motif Conditioning and Scaffold In-painting using Text-Guided Protein Design with ProteoScribe. The left panel shows the wild-type structure of the compact calmodulin (PDB:1PRW) protein, where the grey regions represent the scaffold, and the blue highlights the functional motif. ProteoScribe generates new scaffold designs conditioned on maintaining the functional motif using a text prompt (Appendix B.3). The three designs (Replica 1, 2, and 3) on the right display the predicted scaffold structures (red) generated using ColabFold with MMSeqs2 [72] for sequence alignment and structure prediction. The functional motif (grey) remains consistent across all designs, highlighting the success of the model in preserving motif integrity while generating diverse scaffold structures.

The count statistics for time points *t*_0_[0*M*],*t*_72_[0*M*], and *t*_72_[1*M*], where 0*M* and 1*M* indicate KCl concentration in the media and *t*_*i*_ indicates count frequencies at different time points *i* in hours (Figure S10). These plots display the count distributions for the input population in both trials, population at 72 hours in 0 M KCl and 1 M KCl, providing a baseline for interpreting our results. Figure S11 demonstrates the reproducibility of our experimental results between two independent trials. This figure includes error bars, which were computed using error propagation while assuming Poisson statistics, to illustrate the consistency of our measurements

Figure S12 presents a histogram showing the overall fitness distribution of our designed sequences, and a plot illustrating the percentage of functional rescue achieved by sequences generated from each of the five prompts (Appendix A.5.3). This allows us to assess the general performance of our designs and compare the efficacy of different prompt strategies. A designed sequence is classified as functional if its own uncertainty doesn’t statistically significant overlap with the null allele’s uncertainty. We note that while Prompts 3 and 4 only achieve 0 and 1 *in vivo* functional designs, respectively, the functional designed sequence is the most novel generated among all prompts. With this information, and knowing that the designed sequences were selected based on acceptable TM-score scores, this hints that Prompt 3 and 4 designs sequences might still be foldable and functional sequences, albeit not with a function capable of rescuing *Sho*1 osmosensing in *S. cerevisisae*. Table S9 presents an alignment of the most novel sequence generated from Prompt 4 with the wild-type *Sho*1^*SH*3^ sequence. This table illustrates the poor local alignment and lack of apparent homology between the two sequences, and that this highly divergent sequence, despite its novelty, maintains function while showing minimal similarity to known SH3 domains in BLAST searches.

Table S8 reports *in vitro* biochemical measurements of binding affinity for four designed sequences from Prompt 5 using a tryptophan quenching assay. Variants demonstrate varying affinities for the pbs2 ligand, with Prompt 5, replica 866 showing a significantly lower dissociation constant *K*_*d*_ compared to the wild-type, indicating stronger binding of the designed mutant.

**Figure S10:**
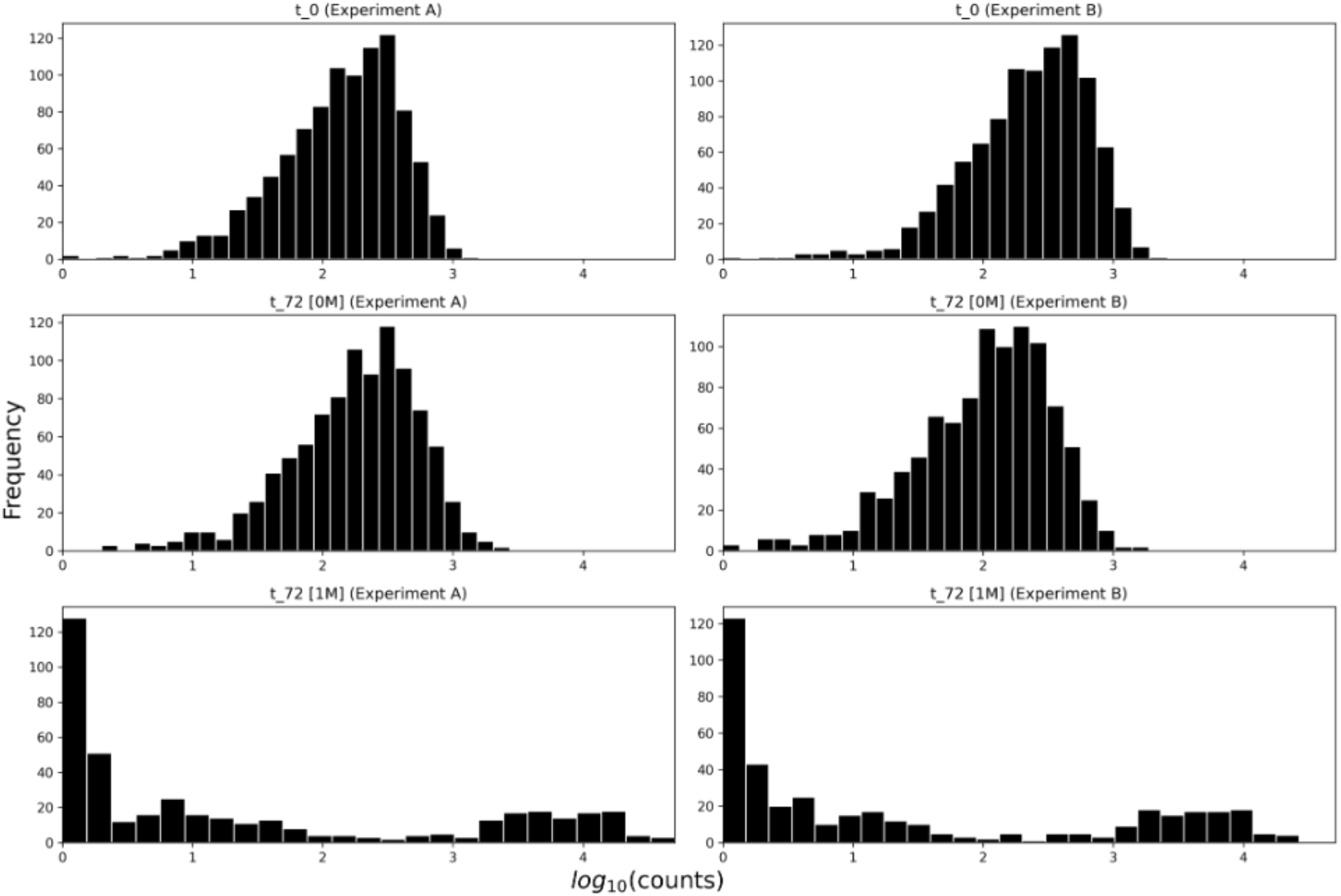
Frequency distribution of sequencing read counts for the *Sho*1^*SH*3^ domain osmosensing selection experiment. Histograms show the log10-transformed read counts at different time points and osmotic conditions: *t*_0_ (initial time point), *t*_72_ [0M] (72 hours, no osmotic stress), and *t*_72_ [1M] (72 hours, 1M KCl osmotic stress) for two independent experiments (A and B). Each row represents a different condition, while columns show experimental replicates. The x-axis displays log10(counts), and the y-axis shows frequency. Data was generated using a MiSeq V2 500 cycle kit, demonstrating reproducibility of sequencing depths and population distributions across samples, time points, and osmotic conditions in the *Sho*1^*SH*3^ domain selection assay.

**Table S8:** Biochemical characterization of wild-type Sho1^SH3^ and designed protein variants from Prompt 5 on pbs2 ligand binding using a tryptophan quenching assay. The table shows the dissociation constant (*K*_*d*_) with mean and standard deviation, relative enrichment (r.e.) scores after 72 hours in selective conditions, and sequence identity to the wild-type Sho1^SH3^. Variants demonstrate varying affinities for the pbs2 ligand, with Prompt 5, replica 866 showing a significantly lower *K*_*d*_ compared to the wild-type, indicating stronger binding. The sequence identities to the wild-type sequence highlight the novelty of the designs while retaining functional binding capabilities.

| Protein | $K_d$ (mean ± std) | r.e. (72 hours) | Seq id to WT (%) |
| --- | --- | --- | --- |
| WT $Sho1^{SH3}$ | $0.9 \pm 0.44$ | 0 | 100 |
| Prompt 5, replica 10 | $1.42 \pm 0.5$ | -0.317 | 45.45 |
| Prompt 5, replica 98 | $2.02 \pm 0.62$ | -0.593 | 51.52 |
| Prompt 5, replica 866 | $0.31 \pm 0.04$ | -0.540 | 47.70 |
| Prompt 5, replica 99 | $1.27 \pm 0.45$ | -2.401 | 46.03 |

**Figure S11:**
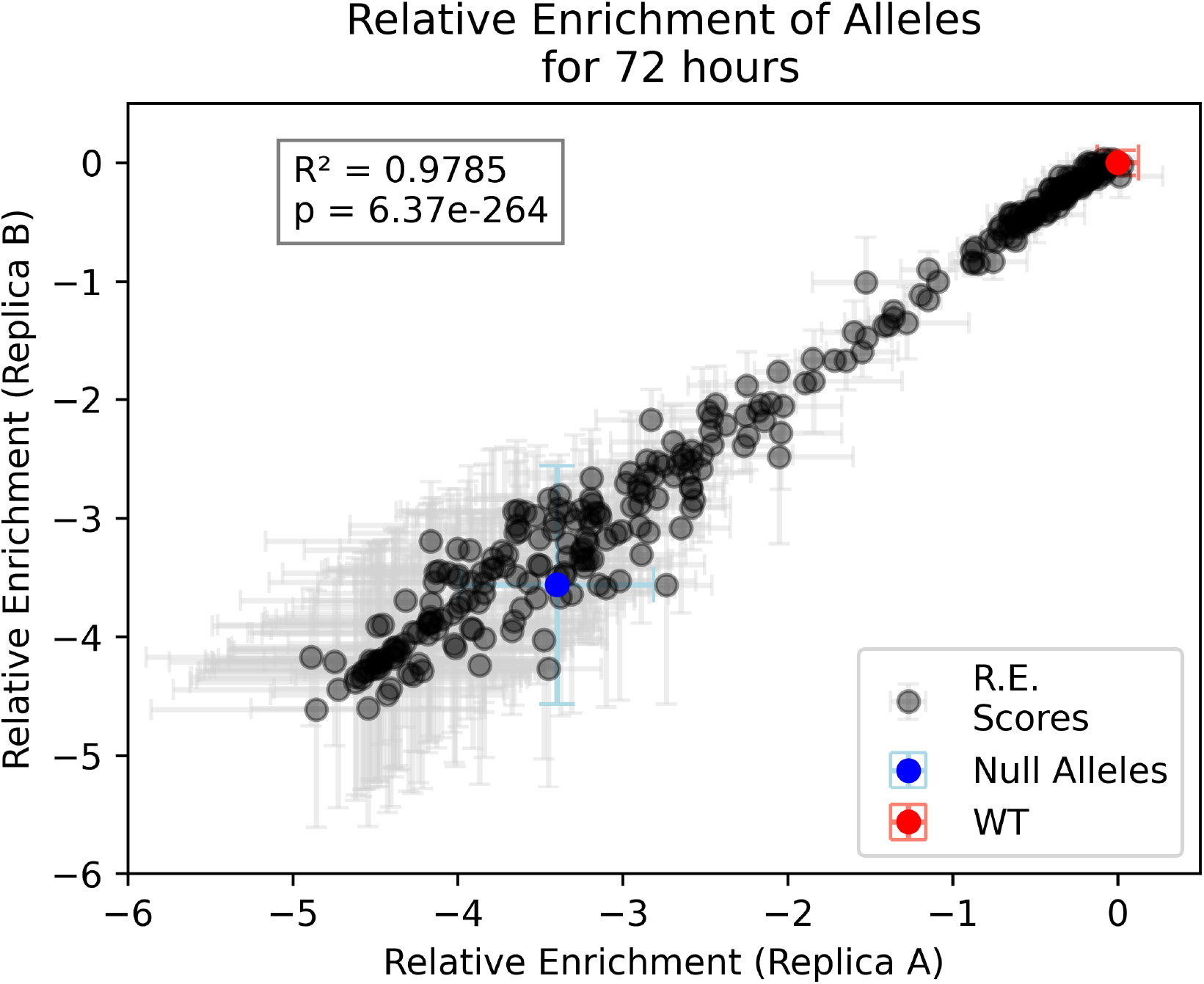
Reproducibility of relative enrichment scores for designed *Sho*1^*SH*3^ domain sequences after 72 hours of selection. The scatter plot compares relative enrichment (r.e.) scores between two experimental replicates (A and B) for text-guided designed *Sho*1^*SH*3^ domain sequences and calibration controls. Gray dots represent individual designed sequences, the blue dot indicates the null allele, and the red dot represents the wild-type (WT) *Sho*1^*SH*3^ domain. Error bars show the uncertainty for each measurement. The high *R*^2^ value (0.9785) and extremely low p-value (6.37e-264) demonstrate strong correlation and statistical significance between replicates, indicating high reproducibility of the selection assay. The plot illustrates the range of fitness effects observed among the designed *Sho*1^*SH*3^ sequences under osmotic stress conditions and serves as a calibration curve for assessing the performance of the designs.

**Table S9:** Comparison of the most novel design sequence against any SH3 domain and the wild-type Sho1^SH3^, along with a search for the closest hits using BLASTp against the NCBI non-redundant database.

| Attribute | Value |
| --- | --- |
| Protein | Prompt 4, replica 403 |
| r.e. [72 hours] (mean ± std) | -1.73 ± 0.10 |
| Sequence identity to any SH3 (%) | 38 |
| Sequence identity to WT (%) | 28 |
| Classification | Functional |
| Closest BLASTp Hit (NCBI) | brain-specific angiogenesis inhibitor 1-associated protein 2-like isoform X2 |

**Figure S12:**
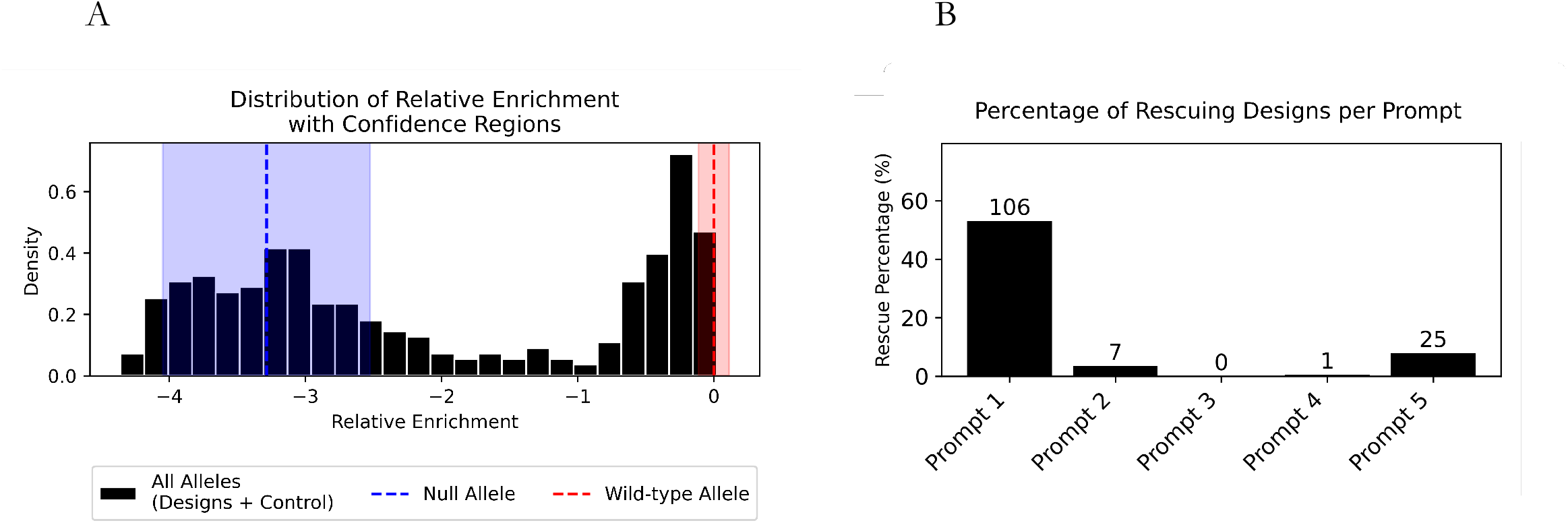
Distribution of relative enrichment scores and rescue efficiency of designed *Sho*1^*SH*3^ domains. (A) Histogram showing the distribution of relative enrichment (r.e.) scores for all sequences, including designed and control alleles. The blue dashed line and shaded region represent the null allele and its confidence interval, while the red dashed line indicates the wild-type allele. (B) Bar plot displaying the percentage of designs that rescue *Sho*1^*SH*3^ function for each of the five prompts (Appendix A.5.3). Rescue is determined by designs having relative enrichment scores significantly higher than the null allele’s confidence region. Numbers above each bar indicate the count of rescuing designs among the 200, 200, 67, 200, and 317 sequences experimentally tested for Prompts 1, 2, 3, 4, and 5, respectively.

